# Kirigami electronics for long-term electrophysiological recording of human neural organoids and assembloids

**DOI:** 10.1101/2023.09.22.559050

**Authors:** Xiao Yang, Csaba Forró, Thomas L. Li, Yuki Miura, Tomasz J. Zaluska, Ching-Ting Tsai, Sabina Kanton, James P. McQueen, Xiaoyu Chen, Valentina Mollo, Francesca Santoro, Sergiu P. Paşca, Bianxiao Cui

## Abstract

Organoids and assembloids have emerged as a promising platform to model aspects of nervous system development. Longterm, minimally-invasive recordings in these multi-cellular systems are essential for developing disease models. Current technologies, such as patch-clamp, penetrating microelectrodes, planar electrode arrays and substrate-attached flexible electrodes, do not, however, allow chronic recording of organoids in suspension, which is necessary to preserve their architecture. Inspired by the art of kirigami, we developed flexible electronics that transition from a 2D pattern to a 3D basket-like configuration to accommodate the long-term culture of organoids in suspension. This platform, named kirigami electronics (KiriE), integrates with and enables chronic recording of cortical organoids while preserving morphology, cytoarchitecture, and cell composition. KiriE can be integrated with optogenetic and pharmacological stimulation and model disease. Moreover, KiriE can capture activity in cortico-striatal assembloids. Moving forward, KiriE could reveal disease phenotypes and activity patterns underlying the assembly of the nervous system.

## Introduction

Self-organizing neural organoids and assembloids derived from human induced pluripotent stem (hiPS) cells hold promise for studying human neurodevelopment including the generation of cell diversity, cell migration, and early circuit assembly^1,2^. One of the goals of these *in vitro* systems is to reveal how electrical activity arises and changes as the nervous system develops and in the context of neuropsychiatric disorders. However, longitudinal tracking of these processes over months is non-trivial as these 3D cultures are often maintained in suspension to self-organize.

Fluorescence imaging methods, such as those using calcium indicators^3^ and voltage sensors^4^, are widely used to measure activity in neuronal populations but limited by their temporal resolution, photobleaching or phototoxicity. Electrophysiological recordings, which are considered the gold standard for monitoring electrical activity, have the potential to monitor neural activity over many months with high temporal resolution. Previous studies monitoring electrical activity in neural organoids using patch-clamp^5,6^, rigid penetrating electrodes^7^ or planar multielectrode arrays (MEAs)^8-11^ are either acute or require slicing the organoids. Recent efforts using buckled MEAs^12^, self-folding shell MEAs^13^, and stretchable mesh MEAs^14^ have enabled electrical recordings from neural organoids without insertion or slicing. These approaches, however, require organoids to be in contact with the substrate, which may interfere with their cytoarchitecture and development.

In this study, we employed the concept of kirigami – the art of paper cutting and 3D folding, to develop an electrical recording platform that can seamlessly integrate with and chronically record from intact neural organoids and assembloids in suspension without interfering with their 3D self-organization and differentiation. These ultra-thin electronic devices, kirigami electronics (KiriE), are geometrically tailored to accommodate neural organoids in suspension, which enables their long-term integration and continuous recording. We demonstrate that KiriE is capable of monitoring spontaneous neural activity in human cortical organoids (hCOs), detecting disease-related phenotypes, and capturing activity in cortico-striatal assembloids.

## Results

### KiriE design, optimization, and device fabrication

To chronically integrate with and record from 3D neural organoids in suspension, we employed the kirigami concept to design an ultra-thin 2D kirigami pattern that spontaneously transforms into a 3D geometry upon release in suspension (**Fig. 1a** and **Supplementary Video 1**). The 3D geometry is a basket with a diameter of 1 cm and 32 microelectrodes (25μm diameter) located in a 1 mm central area. KiriE patterns are nanofabricated with two SU-8 insulation layers encapsulating metal connections for a total thickness of approximately 0.9 μm. In the central area integrating with organoids, SU-8 and metal interconnects are 9 μm and 3 μm wide, respectively.

**Fig. 1.**
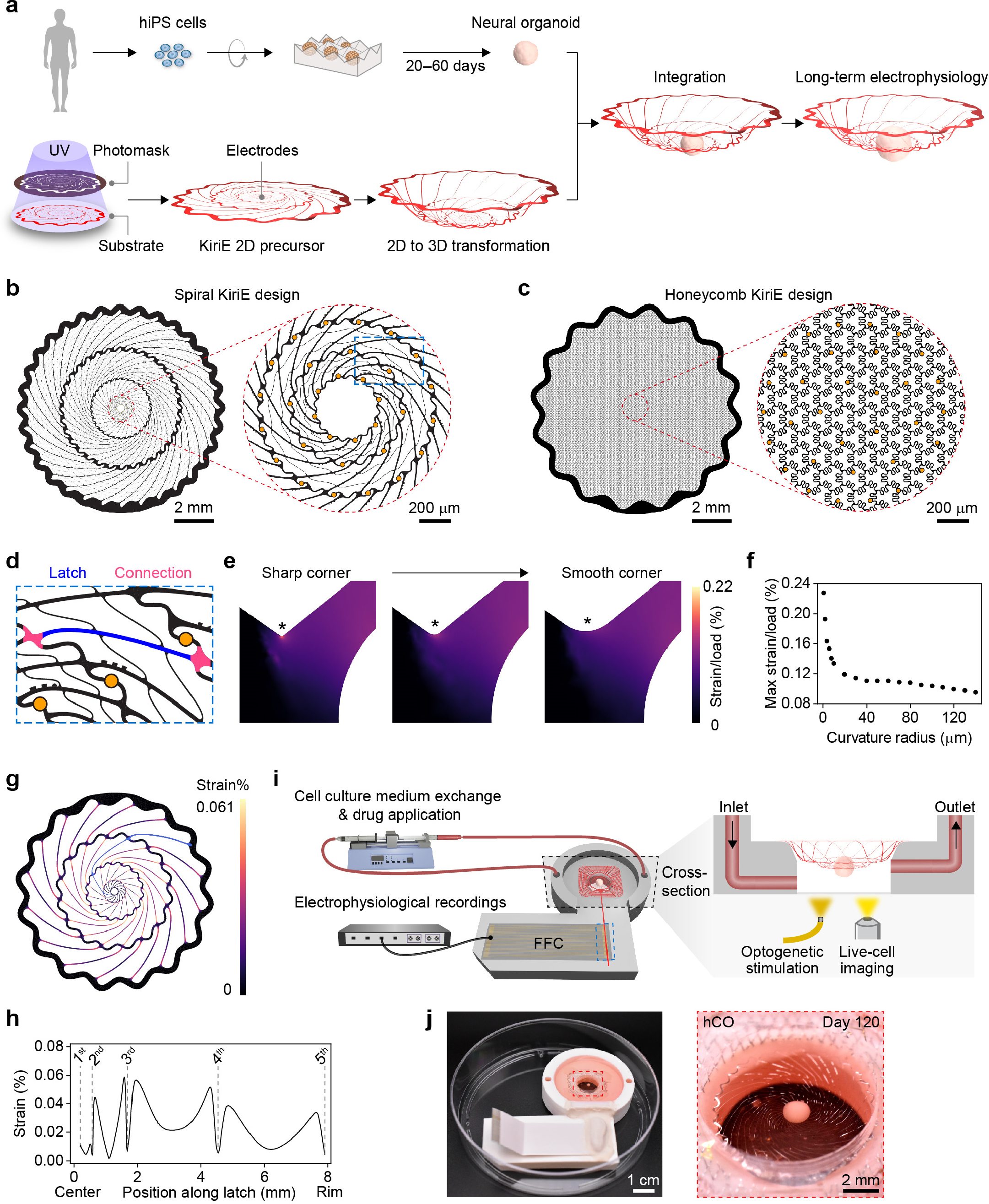
Design, optimization, and fabrication of KiriE for long-term KiriE-hCO integration. **a**, Schematics of the generation of neural organoids from hiPS cells, the concept of vertically deformable KiriE, and the integration of neural organoids with KiriE. **b**, The spiral KiriE design consists of concentric rings connected by spiral latches with 32 electrodes (gold circles) in the central area. **c**, The honeycomb KiriE design is the tiling of serpentine-lined hexagonal units with 32 electrodes in the central area. **d**, Inset of the spiral KiriE (the blue dashed box in **b**) showing a spiral latch (blue) connecting two wavy rings with smooth connections (red). The microelectrodes (gold) and their barcodes are also visible. **e**, FEM simulation shows that the maximum strain accumulates at sharp corners and can be significantly diminished by smoothing the connection (indicated by asterisks). **f**, Quantifications show that rounding off corners (increasing curvature radius) significantly diminishes the strain. **g**, Strain distribution in a simplified spiral KiriE under its own weight and the weight of a 1.2 mm-diameter organoid in the medium. The maximum strain is 0.06%, well below the strain at the elastic limit of SU-8. **h**, The plot of strain along the metal interconnect of a microelectrode (the blue latch in **Fig. 1g**). There is minimal strain between the first and the second rings, where the organoid is located. The vertical dashed lines indicate the positions of five concentric rings. **i**, A schematic of the KiriE platform and its functional modules. FFC, flat flexible cable. **j**, Photographs of a spiral KiriE-hCO assembly. The zoomed-in photograph shows a spherical hCO on a spiral KiriE at day 120 of differentiation.

To design KiriE patterns with substantial deformability and robust mechanical stability that can withstand potential stresses over months of organoid growth, medium exchange and handling, we designed two KiriE patterns – spiral and honeycomb patterns (**Fig. 1b,c**). The spiral pattern consists of concentric rings interconnected by spiral latches, which rotate and vertically extend under a load. The honeycomb pattern is a tiling of serpentine-lined hexagonal units, which expand in size under a load. We developed scripts to create complex KiriE patterns based on a set of input parameters (**Extended Data Fig. 1a–i** and **Methods**) and automatically find the interconnect lines (**Extended Data Fig. 1j–o** and **Methods**). To iteratively simulate and optimize the design, we developed a pipeline combining scripted design with a finite element method (FEM) simulation framework that quantifies deformation under a load and stress accumulations at different locations (**Methods**).

To achieve high deformability and mechanical durability in the spiral design, we identified and iteratively optimized three critical parameters. The first is the shape of connection between the ring and the latch (**Fig. 1d**). FEM simulations show that the strain accumulates at the corner when the ring and the latch are connected at a sharp angle (**Fig. 1e,f**). The maximum strain can be significantly reduced by rounding off the sharp corner and designing wavy rings to reduce horizontal tensions. The second parameter is the shape of the spiral latch, which affects the vertical extension and maximum strain when the latch length, width, and connecting points are kept the same (**Extended Data Fig. 2a–e**). From 150 simulated shapes, we optimized the latch shape for its overall smoothness to reduce local strain accumulation (**Extended Data Fig. 2f,g**). The third parameter is the latch length, which is determined by the location of the connecting point in the inner ring. With the optimized shape smoothness, the vertical deformation depends linearly on the latch length (**Extended Data Fig. 2h**). We chose a latch length that allows sufficient deformation without too much ring rotation during vertical extension that would compromise durability. Our final spiral design had a maximum strain of 0.06% when the pattern is subjected to the weight of KiriE itself and an organoid ranging from 0.2 mm to 1.2 mm in diameter in cell culture medium (**Fig. 1g** and **Extended Data Fig. 2i–k**), which is well below the strain (2–4%) at elastic limit of SU-8^15^. A plot of strain along the metal interconnect (the blue curve in **Fig. 1g**) shows minimal strain between the first and second rings, where the organoid resides (**Fig. 1h**).

To build a highly deformable and mechanically durable honeycomb pattern, we first designed a node that smoothly connects three serpentine lines to avoid sharp angles and local stress accumulations (**Extended Data Fig. 3a**). Then, we designed the serpentine line shape by leveraging prior knowledge of how serpentine deformability is affected by geometric variables^16^. The nodes and serpentines are arranged into a hexagonal unit and tiled within the device area (**Extended Data Fig. 3b–d**). The resulting honeycomb design shows a maximum strain of 0.026% under the weight of KiriE itself and a 1.2-mm-diameter organoid in medium (**Extended Data Fig. 3e,f**). KiriE devices with the optimized patterns were nanofabricated by photolithography (**Extended Data Fig. 3g–l** and **Methods**). Channel-indexing barcodes are incorporated for identification of electrodes (**Extended Data Fig. 3m,n**).

### KiriE enables long-term integration with neural organoids in suspension

To integrate KiriE with organoids, we derived hCOs from hiPS cells as previously described^17,18^ (**Fig. 1a** and **Extended Data Fig. 4a,b**). We designed and 3D-printed a culture chamber that holds KiriE in suspension and is compatible with live-cell imaging, long-term electrophysiology, pharmacological application and optogenetic modulation (**Fig. 1i**). The inlet and outlet channels enable gentle and efficient media perfusion (**Extended Data Fig. 5a–e**). Upon the release of KiriE from the wafer, KiriE is transferred onto the culture chamber with a polydimethylsiloxane (PDMS) supporting ring attached to the area outside the KiriE pattern (**Extended Data Fig. 5f–j**). The contact pads located at the tail of KiriE are aligned with a flat flexible cable (FFC) (**Extended Data Fig. 5k**). Upon drying the tail region, electrical connection is established by direct contact interfacing between KiriE and FFC^19^. In our setup, electrophysiological measurements are performed *in situ*, which can be coupled with the built-in drug perfusion and in-incubator optogenetic stimulation. Moreover, the glass bottom of the chamber allows light access, and therefore enables longitudinal characterization of hCO morphology, live imaging and optogenetic stimulation.

To investigate KiriE-hCO integration, we nanofabricated fluorescent KiriE by covalently incorporating a Rhodamine fluorophore within the SU-8 polymer^20^. We derived hCOs from an hiPS cell line that expresses a green fluorescent protein (CAG::eGFP). We gently pipetted an hCO into the center of KiriE (**Fig. 1j**). Confocal imaging demonstrated that, as expected, both spiral and honeycomb KiriEs are highly deformable (**Fig. 2a** and **Extended Data Fig. 6a**). Fluorescence intensity of CAG::eGFP cells remained relatively constant in relation to the distance from KiriE (**Extended Data Fig. 6b**). After 26 days, KiriE was embedded in hCOs as deep as 300–500 μm (**Extended Data Fig. 6c,d** and **Supplementary Video 2**). Moreover, hCOs on KiriE maintained a spherical morphology similar to hCOs maintained in suspension in ultra-low attachment plates (**Fig. 2b**). We note that only the bottom part of the hCOs was live imaged due to the limited penetration depth. KiriE devices can be cleaned and reused for integration with new organoid cultures (**Extended Data Fig. 5l–p**).

**Fig. 2.**
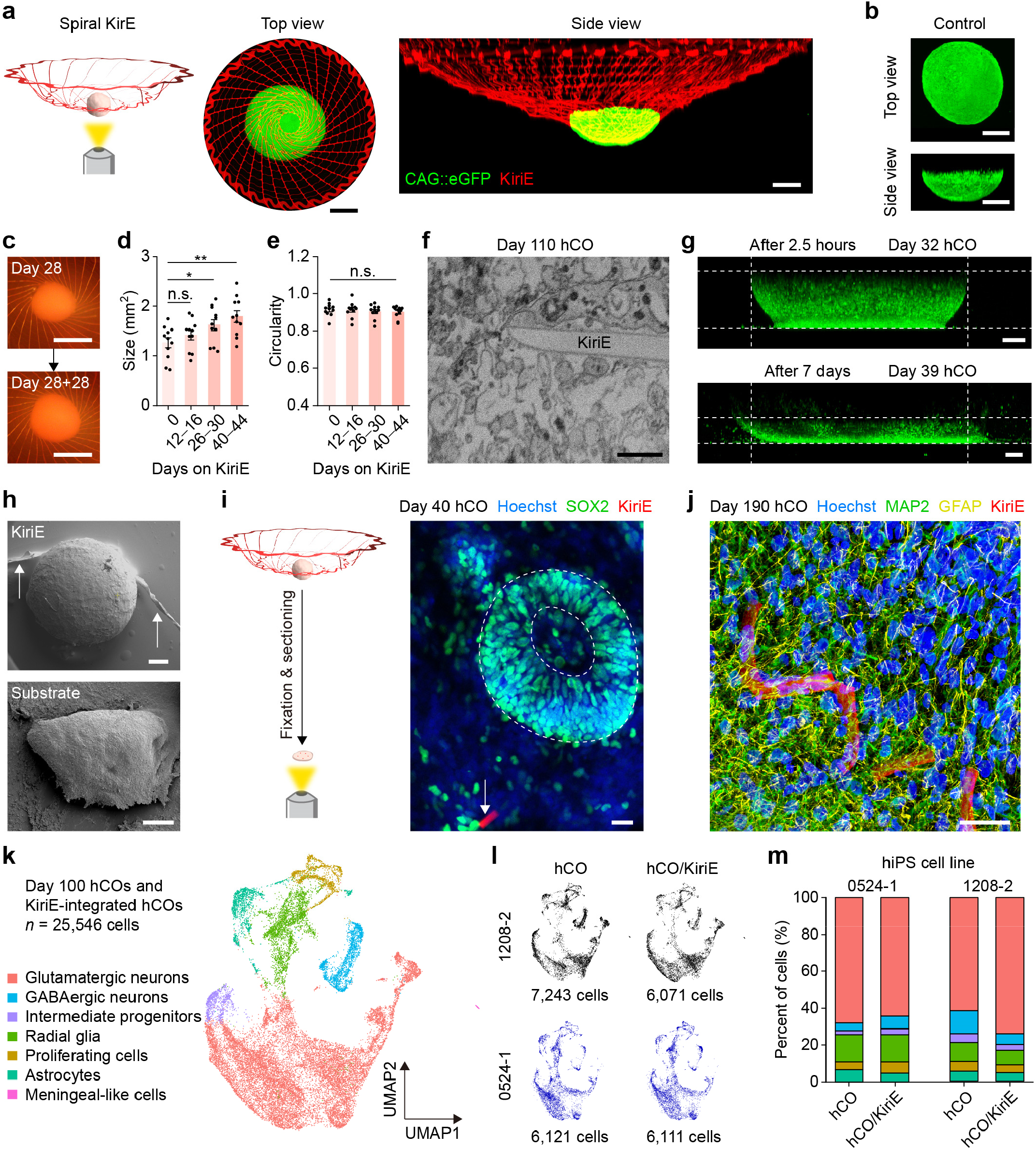
KiriE enables long-term organoid integration in suspension without interfering with hCO development. **a**, The top view and the side view of CAG::eGFP hCOs (green) on the spiral KiriE (red) reconstructed from live-cell confocal imaging. The level of the medium was lowered to better visualize the deformation of KiriE. Scale bars, 500 μm. **b**, The top view and the side view of a CAG::eGFP hCO cultured in suspension (control). Scale bars, 500 μm. **c**, Images of an hCO integrated with a spiral KiriE at different stages of differentiation. These images show that the same hCO continued to grow in size when it was cultured on KiriE. Scale bars, 1 mm. **d**, The plot of size of hCOs as they were cultured on KiriEs over time (*n* = 11 hCOs). One-way ANOVA test, *F*_3,40_ = 4.74, *P* = 6.37 × 10^−3^. Left to right, *P* = 0.314, \**P* = 2.27 × 10^−2^, \*\**P* = 1.09 × 10^−3^. **e**, The plot of circularity of hCOs as they were cultured on KiriEs over time (*n* = 11 hCOs). One-way ANOVA test, *F*_3,40_ = 0.335, *P* = 0.800. Data in **d** and **e** are presented as mean ± s.e.m. **f**, A high-resolution FIB-SEM image showing a KiriE segment in close contact with cells inside an hCO. Scale bar, 2 μm. **g**, Time-lapse live-cell confocal images showing the flattening of a LMNB1-mEGFP hCO after being plated on the adhesive substrate (glass-bottom plate). Additional time-lapse data points are included in **Extended Data Fig. 7a**. Scale bars, 100 μm. **h**, SEM images of an hCO integrated with KiriE for 60 days (top), and an hCO after being cultured on the adhesive substrate for 14 days (bottom). The bundled KiriE structures are indicated by the arrows. Scale bars, 200 μm. **i**, Immunostaining of SOX2^+^ neural progenitors shows the presence of KiriE (red) in or near neural rosettes in KiriE-hCO cryosections at day 40 of differentiation (20 days post-integration). Blue, green, and red colors represent Hoechst, SOX2, and KiriE, respectively. Scale bar, 20 μm. **j**, Immunostaining shows that chronic integration of KiriE did not perturb the spatial distributions of MAP2^+^ neurons or GFAP^+^ glial lineage cells in KiriE-hCO cryosections at day 190 of differentiation (170 days post-integration). Blue, green, yellow, and red colors represent Hoechst, MAP2, GFAP, and KiriE, respectively. Scale bar, 20 μm. **k**, UMAP visualization of scRNA-seq data of day 100 hCOs (*n* = 25,546 cells from *n* = 4 samples of control hCOs and KiriE-integrated hCOs from two hiPS cell lines). hCOs were integrated with KiriE at day 60 of differentiation. **l**, UMAP visualization of scRNA-seq data for control hCOs and KiriE-integrated hCOs color-coded by hiPS cell lines. **m**, Graph showing the percentage of cells belonging to each cluster in each of the *n* = 4 samples.

We continuously monitored hCOs on KiriE, and observed that hCOs cultured on either spiral KiriE (**Fig. 2c**) or honeycomb KiriE (**Extended Data Fig. 6e**) grew in size (**Fig. 2d**) and maintained their spherical morphology over 44 days (**Fig. 2e**). We found that nanofabricated spiral KiriE devices usually have fewer connection defects as their electrode routing lines (∼8 mm) are shorter than those in the honeycomb pattern (∼17 mm). Moreover, the integration of hCO with spiral KiriE with easily identifiable center is more straightforward. Therefore, we primarily focused on spiral KiriE.

To verify the integration of KiriE with hCOs, we employed focused ion beam scanning electron microscopy (FIB-SEM) to examine the KiriE-hCO interface. KiriE-hCO assemblies were fixed, excised from the culture chamber, stained with heavy metal, and then embedded into epoxy resins using an ultrathin resin embedding method (**Methods**). FIB-SEM images showed the deformation of KiriE as it penetrates into the hCO (**Extended Data Fig. 6f**) and, at ∼180 μm depth within hCO, KiriE came in close contact with cells (**Fig. 2f**).

To test the impact of 3D suspended cultures versus attached cultures, we carried out time-lapse live-cell confocal imaging and SEM imaging of hCOs in suspension (control or KiriE) or in contact with an adhesive, rigid substrate. We observed dramatic and time-dependent flattening of hCOs derived from the hiPS cell line LMNB1-mEGFP after contact with an adhesive substrate (**Fig. 2g** and **Extended Data Fig. 7a**). After 28 days of culture, hCOs took a flat shape with reduced thickness of 100–200 μm (**Extended Data Figure 7a**). Moreover, SEM images showed that hCOs maintained their spherical shape after 60 days on KiriE, while hCOs plated on the adhesive substrate had flattened after 14 days (**Fig. 2h**). These suggest that maintaining cultures in suspension is critical to preserving 3D morphology. Moreover, SYTOX™ deep red dead cell stain indicated a lower cell viability in hCOs cultured on the adhesive substrate compared to hCOs cultured in suspension (control) or on KiriE after 40–45 days of culture (**Extended Data Fig. 7b,c**). Similarly, the expression of the apoptosis marker cleaved caspase-3 (c-Cas3) in day 91–102 hCOs was higher in hCOs on the adhesive substrate than in hCOs kept in suspension (control) or on KiriE (**Extended Fig. 7d,e**). These results indicate that maintaining 3D cultures in suspension is associated with improved cell health.

### KiriE integration does not interfere with hCO development

To examine whether KiriE affects the cytoarchitectural organization of hCOs, we placed hCOs on KiriE at day 20 of differentiation and examined them up to day 190. As previously described^17,18,21^, we found SOX2^+^ neural progenitors organized in the ventricular zones (**Fig. 2i** and **Extended Data Fig. 6g,h**) as well as expression of neuronal (MAP2) and glial cell lineage (GFAP) markers (**Fig. 2j**). Fluorescence analysis as a function of distances from KiriE showed relatively uniform distribution of SOX2^+^, MAP2^+^, and GFAP^+^ cells at the KiriE-hCO interface (**Extended Data Fig. 7f,g**).

To investigate whether KiriE integration interferes with corticogenesis, we performed single cell RNA sequencing (scRNA-seq) of KiriE-integrated hCOs and control hCOs maintained in ultra-low attachment plates at day 100 (*n* = 25,546 cells from *n* = 4 samples from two hiPS cell lines). Uniform Manifold Approximation and Projection (UMAP) dimensionality reduction revealed seven major cell clusters (**Fig. 2k** and **Extended Data Fig. 4c–g**), including glutamatergic neurons expressing *SLC17A7* and *SLC17A6* (66.59%), radial glia expressing *MOXD1* (11.85%), proliferating cells expressing *TOP2A* (5.00%), intermediate progenitors expressing *EOMES* (3.46%), and astroglial lineage cells expressing *AQP4* (5.26%). Next, we compared cell composition between KiriE-integrated hCOs and control hCOs (**Fig. 2l**) and found a similar distribution of cell types for both hiPS cell lines (**Fig. 2m**). Pearson analysis of normalized gene expression showed high levels of correlation between hCOs and KiriE-integrated hCOs (**Extended Data Fig. 4g**). These experiments suggest that incorporation of KiriE into hCO does not interfere with corticogenesis.

### Electrophysiological recordings and modulation of activity in KiriE-hCO

We next performed recordings of spontaneous neuronal activity in intact hCOs from day 75 to 179 of hCO differentiation (**Fig. 3a** and **Extended Data Fig. 8a**). We did not detect spontaneous activity at day 75 but detected significant activity in the same hCO after day 96. Analysis of single-unit activity across channels showed that an individual electrode can detect as many as four neurons (**Fig. 3b** and **Extended Data Fig. 8b–f**). The action potentials exhibited characteristic waveforms with spike amplitudes primarily in the range of 30–140 μV, which remained stable across days (**Extended Data Fig. 8g**). The majority of single-unit spikes were stable for days, while some signals emerge and disappear over time. Longitudinal measurements of multiple KiriE-hCO assemblies showed that neural activity arises around day 100 ± 5 of differentiation (**Fig. 3c**). The electrode impedance in the KiriE-hCO assemblies remained relatively stable at ∼0.3 MΩ (**Extended Data Fig. 8h**). Moreover, firing rates of day 116–125 hCOs measured by KiriE are similar to those measured by acutely inserted silicon shank electrodes (**Extended Data Fig. 8i**).

**Fig. 3.**
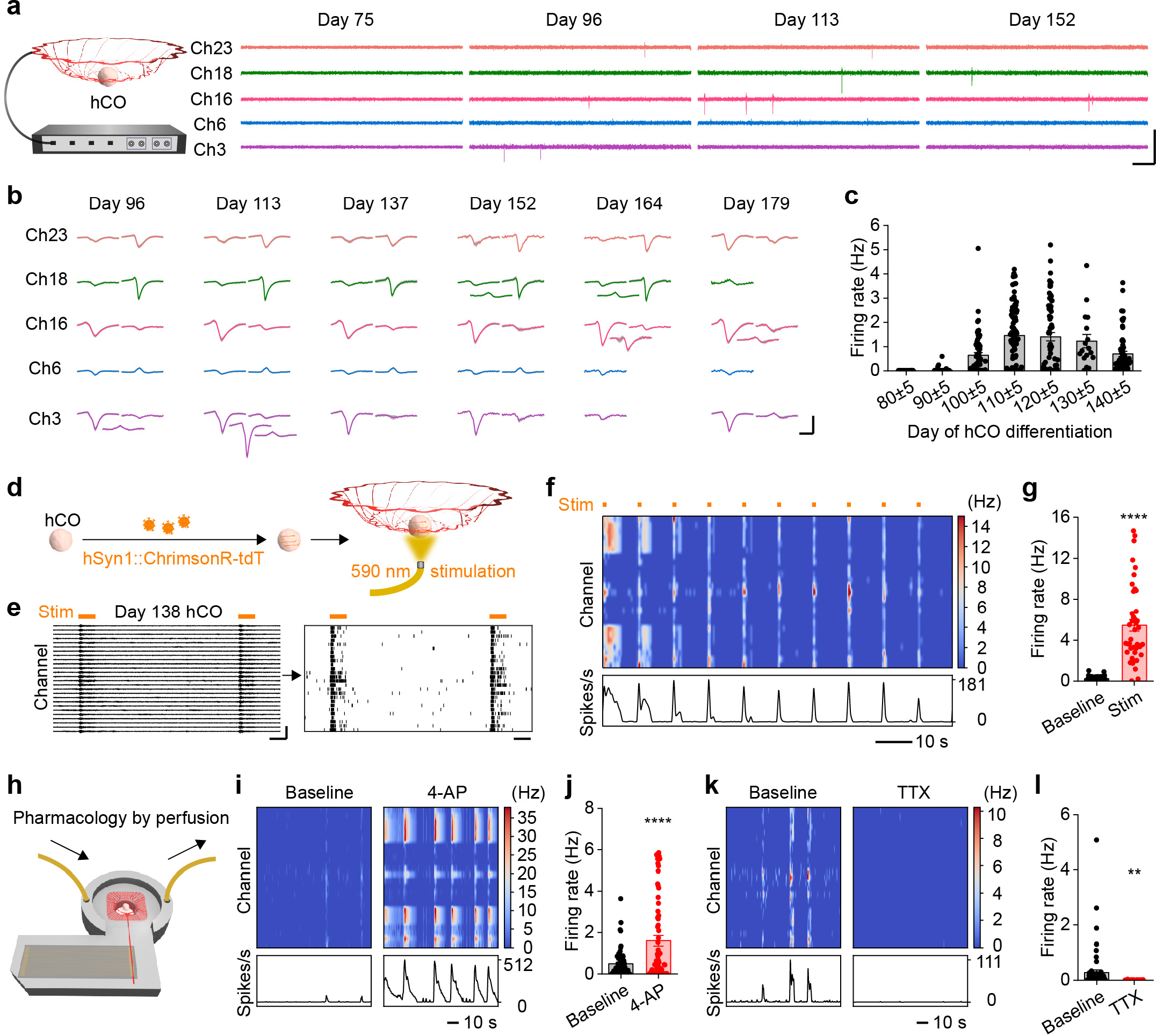
KiriE enables long-term single-unit recordings from intact hCOs. **a**, Representative electrophysiological recordings of spontaneous electrical activity of an hCO at days 75, 96, 113, and 152 of differentiation, respectively. Scale bars, 100 ms (lateral) and 200 μV (vertical). **b**, Average spike waveforms of single unit clusters from 5 representative KiriE channels on days 96, 113, 137, 152, 164, and 179. Scale bars, 1 ms (lateral) and 100 μV (vertical). **c**, Quantification of hCO firing rates as a time course (*n* = 81 channels from four hCOs). **d**, Schematic of the optogenetic stimulation. 590 nm light is delivered to the hCO from the bottom through a fiber optic cannula inside the incubator. **e**, A representative electrophysiological recording and the corresponding raster plot of the spiking activity during the optogenetic stimulation of a day 138 hCO. The orange horizontal lines indicate the 1 s duration of 590 nm light stimulation. Scale bars, 1 s (lateral) and 200 μV (vertical). **f**, The firing rate heatmap during 10 pulses of optogenetic stimulation (*n* = 26 channels). The orange lines indicate the durations of 590 nm light stimulation. **g**, The quantification of firing rates during the light-off and the light-on optogenetic stimulation phases (*n* = 43 channels from two hCOs at day 130–140 from two hiPS cell lines analyzed over 32 light pulses). 0.24 ± 0.03 Hz for light-off periods. 5.44 ± 0.57 Hz for light-on periods. \*\*\*\**P* < 1 × 10^−4^; two-tailed *t*-test. **h**, A schematic drawing of the drug application through the builtin perfusion system in the multimodal culture chamber. **i**, The firing rate heatmaps show the single-unit activity of a day 138 hCO before and after the application of 4-AP (50 μM) (*n* = 25 channels). **j**, The quantification of firing rates before and after the application of 4-AP (*n* = 67 channels from three hCOs). \*\*\*\**P* < 1 × 10^−4^; two-tailed *t*-test. **k**, Firing rate heatmap showing the single-unit activity before and after the application of TTX (1 μM) (*n* = 25 channels). **l**, Quantification of firing rates of baseline and TTX conditions (*n* = 67 channels from three hCOs). \*\**P* = 1.83 × 10^−3^; two-tailed *t*-test. Data are presented as mean ± s.e.m. (**c,g,j,l**).

We manipulated activity in hCO and verified if KiriE can reliably detect neural responses. We infected hCOs with an adeno-associated virus (AAV) expressing hSyn1::ChrimsonRtdTomato –an opsin that activates neurons in response to 590 nm light^22^ (**Fig. 3d**). We delivered light from the bottom of the chamber using an optical fiber (**Extended Data Fig. 8j**) and reliably detected light-evoked responses in hCOs (**Fig. 3e,f**). Quantification of the average firing rates over all channels shows that firing rates during light-on periods were ∼20 times higher than during light-off periods (**Fig. 3g**).

Next, we tested the ability of the KiriE-hCO to detect responses to pharmacological manipulation using the built-in perfusion system (**Fig. 3h**). The application of the potassium channel blocker 4-aminopyridine (4-AP) led to a threefold increase in firing rate (from 0.49 ± 0.08 Hz to 1.60 ± 0.25 Hz; **Fig. 3i,j**). In contrast, application of tetrodotoxin (TTX) – a sodium channel blocker, almost completely abolished spontaneous activity (**Fig. 3k,l**), which was reversible upon washout (**Extended Data Fig. 8k**).These results show that KiriE can record spontaneous activity in hCOs over time and can be integrated with optogenetic and pharmacological perturbations.

### Detecting disease-related phenotypes in KiriE-hCO

To verify the ability of KiriE-hCO to identify disease phenotypes, we integrated KiriE with hCO derived from an hiPS cell line carrying a heterozygous loss of the *DGCR8* gene (**Fig. 4a**). We previously showed that dissociated cortical neurons derived from this mutant line (or from patients carrying the 22q11.2 deletion that encompasses *DGCR8*) display increased spontaneous activity using calcium imaging and patch-clamp^23^. Electrophysiological recordings showed that the spontaneous firing rate of DGCR8^+/-^ KiriE-hCOs at day 133–141 was approximately three times of the control KiriEhCOs at day 137–138 (**Fig. 4b,c** and **Extended Data Fig. 9a**). About 30% of the channels probing DGCR8^+/-^ hCOs showed firing rates greater than 1 Hz (in contrast to 2% in control hCOs), and some were as high as ∼7 Hz. Therefore, KiriE can capture disease-related phenotypes in intact hCOs without dissociation or slicing.

**Fig. 4.**
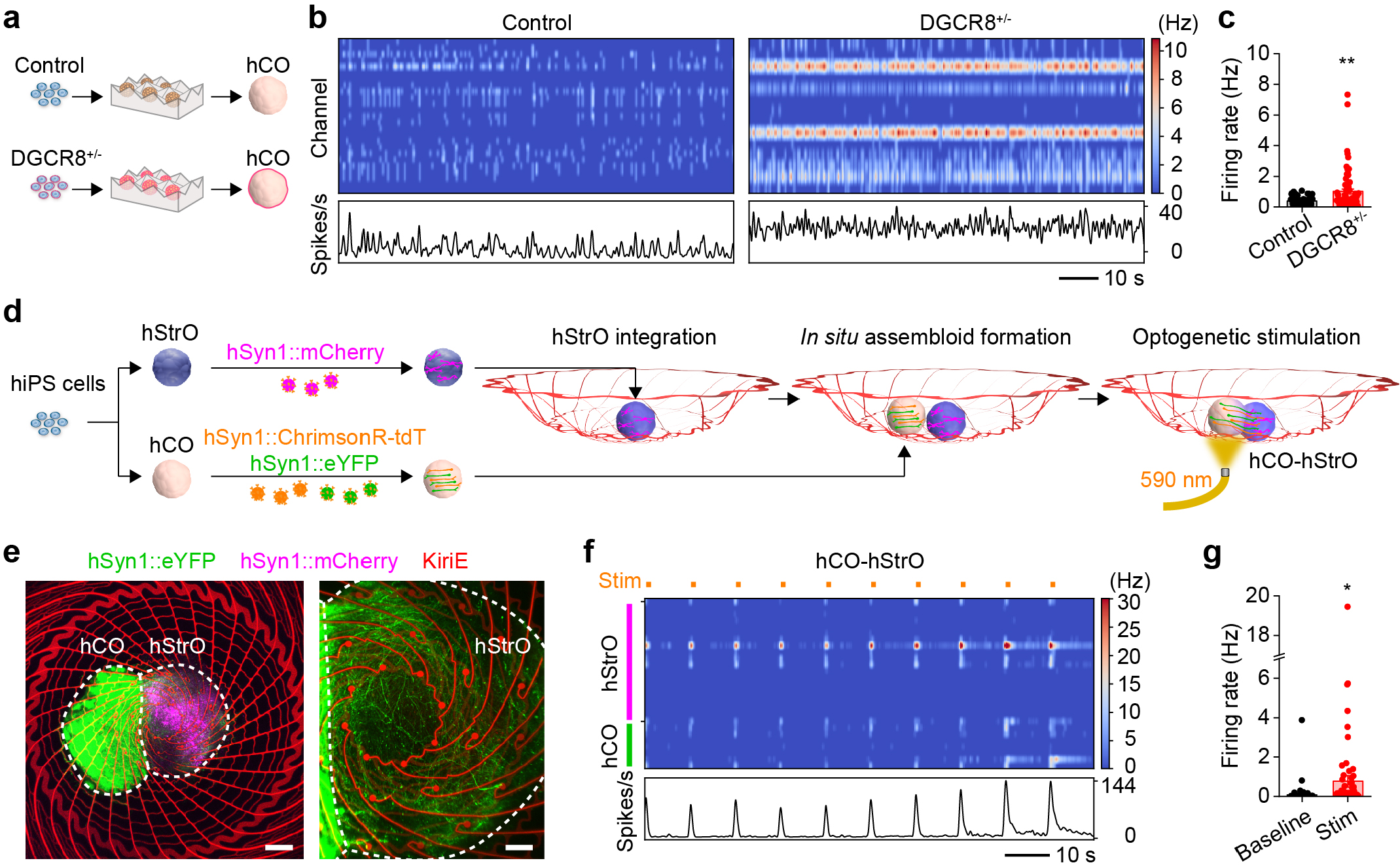
KiriE recording captures disease-related phenotypes and neural activity in cortico-striatal assembloids. **a**, Schematic illustrating hCOs differentiated from DGCR8^+/-^ hiPS cell line and the isogenic control hiPS cell line. **b**, Representative firing rate heatmaps of spontaneous activity in a control hCO (day 137; *n* = 26 channels) and a DGCR8^+/-^ hCO (day 141; *n* = 15 channels). **c**, Quantifications of firing rates of control hCOs (day 137–138; 0.38 ± Hz; *n* = 51 channels from two hCOs) and DGCR8^+/-^ hCOs (day 133–141; 1.01 ± 0.16 Hz; *n* = 70 channels from four hCOs). \*\**P* = 1.2 × 10^−3^; twotailed *t*-test. **d**, The experimental design of forming a cortico-striatal assembloid on a KiriE and interrogating their circuit connectivity by optogenetic stimulation and KiriE recording. **e**, Live-cell confocal fluorescence images of a representative cortico-striatal assembloid. Image on the left shows both hCO and hStrO are in contact with some electrodes in the center of KiriE. Image on the right shows a zoomed-in image of the hStrO. The green channel clearly shows the eYFP^+^ hCO projections into hStrO. Scale bars, 300 μm (left) and 100 μm (right). **f**, The firing rate heatmap of neural activity of the corticostriatal assembloid at day 167 of differentiation (*n* = 20 channels in the hStrO and *n* = 7 channels in the hCO). Upon optogenetic stimulation of the ChrimsonR-expressing hCO, both hCO and hStrO show enhanced activity. The orange lines indicate the pulse durations of 590 nm light stimulation. **g**, The quantification of firing rates of hStrOs in the cortico-striatal assembloids during the light-off (baseline) and the light-on (stimulation) phases (*n* = 74 channels from three cortico-striatal assembloids analyzed over 18–60 light pulses). 0.08 ± 0.05 Hz for light-off periods. 0.77 ± 0.29 Hz for light-on periods. \**P* = 2.04 × 10^−2^; two-tailed *t*-test. Data are presented as mean ± s.e.m. (**c,g**).

### Probing functional neural circuits in cortico-striatal assembloids

We previously showed that hCOs can be integrated with human striatal organoids (hStrOs) to form assembloids and study cortico-striatal circuits^24^. The basket-like geometry of KiriE provides an ideal platform to support the *in situ* fusion of organoids into assembloids and long-term monitoring of electrical activity.

To test this, we first differentiated hStrOs from hiPS cells and integrated them with KiriE using the same procedure for hCOs. Immunofluorescence staining of MAP2^+^ neurons and GFAP^+^ glial lineage cells in KiriE-hStrO (**Extended Data Fig. 9b**) demonstrated a seamlessly integrated neural interface. We also successfully obtained single-unit recordings of KiriEhStrOs that were stable across days (**Extended Data Fig. 9c,d**).

We then separately derived hCOs and hStrOs, and infected hStrOs with AAV-hSyn1::mCherry for fluorescence visualization, and infected hCOs with AAV-hSyn1::eYFP and AAV-hSyn1::ChrimsonR-tdTomato for fluorescence visualization and optogenetic stimulation (**Fig. 4d**). To build a cortico-striatal assembloid on KiriE, we first placed an hStrO at day 80–82 of differentiation in the center of KiriE. Two days later, and after the hStrO had adhered to the electrode area, we placed an hCO at the same stage adjacent to the hStrO to assemble.

Projections of eYFP^+^ hCO neurons into hStrO were found in hCO-hStrO-KiriE assembloids as early as ∼30 days postassembly (**Extended Data Fig. 9e,f**); however, minimal projections of mCherry^+^ hStrO neurons into hCO were found (**Extended Data Fig. 9g**), which is a feature consistent with the unilateral cortico-striatal connectivity *in vivo*^25^. We then leveraged the channel-indexing barcodes to identify the subset of KiriE electrodes embedded in hStrOs or in hCOs. In the assembloid shown in **Fig. 4e**, we found 20 electrodes embedded in the hStrO and 7 electrodes in the hCO (top part and bottom part in **Fig. 4f** and **Extended Data Fig. 9h**). Since hCO neurons, but not hStrO neurons, expressed ChrimsonR, we optogenetically activated hCO neurons and probed the evoked responses. We detected increased activity in hCO neurons upon optogenetic stimulation and a corresponding increase in neuronal activity in hStrO neurons (**Fig. 4f,g**). To explore hCO-hStrO connectivity over time, we longitudinally monitored the activity from 3 weeks post-assembly onward and observed light-induced response only after 7 weeks (**Extended Data Fig. 9i**). Together, these results demonstrate that the KiriE-assembloids can probe connectivity across functionally integrated organoids.

## Discussion

We developed KiriE – a kirigami-inspired electrophysiology platform that is flexible, fully suspended in liquid, and deformable in 3D. Live-cell imaging, immunohistochemistry, and single cell transcriptomics suggested that KiriE integration does not perturb the morphology or cell composition of organoids. KiriE can chronically record from intact neural organoids over several months and can detect electrophysiological phenotypes associated with disease. Furthermore, KiriE can be used to monitor the circuit connectivity in assembloids *in vitro*.

The KiriE platform has several advantages compared to prior technologies^5-14,26^. First, the 3D basket-like geometry of KiriE is designed for stable integration with intact neural organoids in suspension without the need for insertion, slicing or contacting the substrate. This strategy helps preserve the selforganization of 3D cultures with minimal perturbation. Second, the multimodal culture platform enables long-term medium perfusion and *in situ* assays including longitudinal morphology monitoring, live-cell imaging, electrophysiology, and optogenetic and pharmacological modulation, without the need to transfer across setups that could perturb development or induce cellular stress. Finally, KiriE detects disease-associated electrophysiological phenotypes and network connectivity in assembloids integrated *in situ*, which could be applicable to other multi-cellular systems such as cardiac organoids and cortico-spinal-muscle assembloids. There are a number of limitations of the KiriE platform. For instance, simulating and optimizing the mechanical behavior of KiriE is computationally intensive, thus posing a potential challenge to design more complex geometric patterns. Furthermore, the process of fabricating and assembling KiriE devices requires advanced expertise. Future developments of this platform could incorporate pH, oxygen, and neurotransmitter sensors, or micro-LEDs for localized optogenetic activation^27,28^. It is also conceivable to design KiriE into complex 3D geometries such as interconnected baskets for probing neural transmission in linear or loop assembloids.

## Methods

### Script-assisted design of KiriE

The designs were generated via the Python package *PHIDL*^29^. Polygons were defined using Boolean operators and the GSD-II output is compatible with photolithography (**Extended Data Fig. 1a**). Numerical libraries of Python were leveraged to help generate complex geometries. The latches of the spiral design were generated via non-uniform rational basis spline (NURBS) curves (**Extended Data Fig. 1b–g**). In brief, two polygons are connected by a latch at user-defined points on each polygon. The normal vectors to each polygon’s outline at the chosen points are calculated. The normal vectors’ magnitude and angle can be adjusted. The adjusted vectors define two control points. In addition to the beginning and end points of the latch, these additional two points allow to generate a 4-point NURBS curve which outlines the general shape of the latch. Normal vectors to the polygons’ outline were used to ensure that the latch connects to the polygons without any mechanically unfavorable kinks. Furthermore, NURBS curves are at least twice continuously derivable, ensuring smoothness of the outline, which is important for mechanical stability. The final layout is built by a generator script that takes parameters as input such as shape controls of the latch, number of rings, latches and electrodes (**Extended Data Fig. 1h**).

For the purpose of automatically routing the electrode lines to the contact pads, Device and Polygon class within *PHIDL* were made where each polygon (latch, electrode, pad, etc) is endowed with extra node-points, and implicit paths within the polygons (**Extended Data Fig. 1j,k**). Once all the polygons are placed, the nodes are converted to nodes in a graph object (*NetworkX* package). The nodes were connected by an edge within this graph if they are close to each other. All the nodes corresponding to electrodes are connected to a common, virtual ‘source’ node, while all the pads are connected to a common, virtual ‘drain’ node. Edmonds-Karp flow algorithm (*NetworkX*) is used to find the node-disjoint path between the source and the drain, which translates to finding non-intersecting (i.e, not short-circuited) paths between electrodes and pads. The graph therefore gives a pathing through polygon-nodes from electrode to pad. See **Extended Data Fig. 2l–o** for details.

### Mechanical analysis of KiriE

The designs generated during the script-assisted design are a collection of polygons. By using *pygmsh*, a Python package bundling the Gmsh finite-element mesh generator^30^, the designs were turned into a triangulated geometry adapted to finite element analysis (**Extended Data Fig. 1i**). Technically, this is done by a Boolean union of domains, where each polygon is treated as a closed-loop domain. The vertical dimension of KiriE (its thickness) is about 10,000 times smaller than its lateral dimensions. Therefore, a shell formulation was employed, whereby the geometry was treated as a 2D surface embedded in 3D space and the pattern thickness implicitly integrated into the equations. This allows for much better triangular meshing of the geometry, with about 80,000 triangles. Non-linear Naghdi-shell formulation was employed to simulate the deformation of KiriE, as it is suited for large displacement simulation of shell mechanics. The weak formulation of the mechanical equations and the choice of functional space were inspired by a previous work using Python’s *FEniCS* package (*FEniCS-Shells*)^31^. *FeniCS* is an open-source finite element method solver package^32^.

The deformation of KiriE was simulated when it is subjected to gravitational loading plus the weight of a 1.2 mm-diameter organoid in medium. The bending, shear, and membrane strain tensors were retrieved to verify whether the compounded strain was within the elastic limit of SU-8.

The general limitation of non-linear shell theories used to simulate KiriE’s mechanical behavior is that the thickness of the material needs to be much smaller than the radius of the curvature in the deformed pattern. This is indeed the case for KiriE (≪ 1%), but this optimization will not apply for devices with thick meshes. Furthermore, the optimization procedure is computationally intensive and thus limits the complexity of the features that can be optimized. Notably, triangle units in the simulations need to be sufficiently small to capture details (such as the connection points), but the execution time scales as N^2^ (N is the number of triangles). Therefore, patterns consisting of fewer than 100,000 triangles can be processed efficiently. While this is already sufficient for our current KiriE design, it poses a potential limit for more complicated patterns.

### Fabrication of KiriE

The fabrication of KiriE is achieved by multi-layer photolithography (PL) as previously described^19,20^ with minor optimizations. The key fabrication steps are as follows: (1) A 80-nm thick Ni sacrificial layer was evaporated (AJA International Inc) onto a 100 mm Si wafer (p-type 0.1-0.9 Ω cm, Pure Wafer). (2) The Si wafer was cleaned with oxygen plasma (100 W, 2 min; Technics PE II-A) immediately before being spin-coated with LOR 3A (Kayaku Advanced Materials) at 1000 rpm and baked at 180 °C for 3 min. Positive photoresist (S1805, Kayaku Advanced Materials) was spin-coated on the Si wafer at 4000 rpm and baked at 115 °C for 3 min. The positive photoresist was patterned by photolithography (PL) with a mask aligner (SUSS MA6 mask aligner, SUSS MicroTec) and developed (MF-26A developer) for 90 s. (3) 5 nm Ti and 100 nm Au were sequentially deposited by electron-beam evaporation (AJA International Inc), followed by a lift-off step (MF-26A) for the Au Input/Output (I/O) pads. (4) Negative photoresist SU-8 2000.5 (Kayaku Advanced Materials) was diluted in cyclopentanone (Sigma-Aldrich) at a 4:1 ratio, spin coated on the Si wafer at 4000 rpm, pre-baked sequentially at 65 °C for 1 min and 95 °C for 3 min, and patterned by PL. The Si wafer was postbaked sequentially at 65 °C for 1 min and 95 °C for 3 min after PL exposure. (5) The negative photoresist was developed (SU-8 Developer, Kayaku Advanced Materials) for 2 min, rinsed with isopropanol, dried and hard baked at 180 °C for 1 h. (6) Negative photoresist SU-8 2000.5 was mixed with Lissamine rhodamine B ethylenediamine (RhBen; ∼10 μg/ml; Thermo Fisher Scientific) and placed in the dark at room temperature (RT) for at least 3 days to afford stable fluorescence labeling^20^. PL using the completely dissolved SU8/RhBen solution was performed by repeating steps 4 and 5 at 3000 rpm to pattern the bottom SU-8 layer. (7) Steps 2 and 3 were repeated for PL patterning of the interconnect layer consisting of 5 nm Ti, 100 nm Au and 50 nm Pt. (8) Step 6 was repeated for PL patterning of the top SU-8 layer as the insulating layer of the interconnect lines. The Si wafer was hard baked at 195 °C for 1.5 h to allow interdiffusion of the bottom and top SU-8 layers.

### Design and production of multifunctional culture platform

3D design of the multifunctional culture platform was performed using Autodesk 3ds Max (Autodesk Inc). Culture platforms were produced using the Original Prusa i3 MK3S+ 3D printer (Prusa Research) with a biocompatible thermoplastic polyester polylactic acid (PLA) filament (Prusa Research). For 3D printing, structures are 100% filled with randomized seams.

### Assembly and electrical connection of KiriE on the culture chamber

The process of assembling KiriE on the culture chamber is illustrated in **Extended Data Fig. 5f–j**. PDMS at a 10:1 ratio (w/w; base:curing agent; Sylgard 184 silicone elastomer, VWR) was cast into a 3D printed square shape mold and cured at 55 °C for 24 h to make PDMS support. The Si wafer with KiriE was treated with oxygen plasma (50 W, 30 s; Technics PE II-A) and then the cured PDMS support was glued on the KiriE using uncured PDMS and the cured at 100 °C for 1 h. Then the Si wafer with PDMS support on top was transferred to a Ni etchant solution comprising 40% FeCl_3_:39% HCl:H_2_O = 1:1:10 for 30 min. Released KiriE with PDMS support were rinsed with 70% isopropanol (IPA) and deionized (DI) water for 3 times. KiriE with PDMS support remains at the liquid-air interface in the DI water until transferred to the 3D culture chamber.

The preparation of the 3D culture chamber includes gluing a #1.5 glass coverslip (VWR) at the bottom and an FFC (Digi-Key) on the top surface. The glass coverslip was glued to the bottom of the culture chamber using PDMS, and cured at 55 °C for 24 h. An FFC was mounted onto the 3D culture chamber with Metabond cement (Metabond quick luting cement; Parkell). The whole chamber was treated for 1 min by oxygen plasma (Basic Plasma Cleaner, PDC-32G, Harrick Plasma) and sterilized in 70% ethanol.

To assemble KiriE on the 3D culture chamber, KiriE was transferred to and aligned with the 3D culture chamber. The culture chamber was initially immersed under water to avoid accidental tear of KiriE. Then, the liquid level was slowly reduced. Electrical connection of the KiriE direct contact interfacing pads to the FFC was performed using a previously described method^19^. In brief, the interfacing pads of KiriE were aligned with the FFC leads using a transfer pipette. Aligned interfacing pads were fully dried by surgical spears (Braintree Scientific) and fixed in place using epoxy adhesive (5 minute epoxy, Devcon). The interface was covered with Metabond cement. Metabond cement was also used to fix the PDMS support of KiriE to the culture chamber. A 3D printed bar was glued to the opening of the circular chamber with Metabond cement to seal the chamber. The culture chamber was housed individually in the 10 cm tissue culture dish.

### Characterization and maintenance of hiPS cells

hiPS cell lines used in this study were validated using standardized methods as previously described^18^. Cultures were regularly tested to ensure they are free of mycoplasma. A total of six control hiPS cell lines derived from fibroblasts from six healthy individuals were used for experiments. DGCR8^+/-^ hiPS cell line was generated as previously described^23^. Approval for this study was obtained from the Stanford IRB panel, and informed consent was obtained from all participants.

### Generation of CAG::eGFP hiPS cell line

CAG::eGFP hiPS cell line was generated as previously described^33^. The parental hiPS cell line (8858-3) was maintained in 6-well plates using StemFlex medium (Life Technologies, A3349401). Cas9, gRNA and donor plasmids were obtained from Addgene (plasmids #42230, #41818 and #52344, respectively). For nucleofection, 3 μg Cas9, 1 μg gRNA and 1 μg donor plasmids were used. On the day of nucleofection, hiPS cells were washed with DPBS and incubated with 1 ml Accutase at 37 °C for 10 min, after which 9 ml Essential 8 medium (Thermo Fisher Scientific, A1517001) was added to the well for resuspension. After cell counting, the single cell suspension containing 3 × 10^6^ cells was centrifuged. The cell pellet was used for nucleofection using the P3 Primary Cell 4D-Nucleofector X Kit L (Lonza, V4XP-3024), a 4D-nucleofector core unit and X unit (Lonza) using the nucleofection program DC100. After nucleofection, the cells were immediately seeded into a well of a 6-well plate that was pre-coated with vitronectin and contained pre-warmed Essential 8 medium supplemented with the ROCK inhibitor Y-27632 (10 μM; Selleck Chemicals, S1049). While the nucleofected cells recovered to reach 70%–80% confluency, 1 μg ml^−1^ of puromycin was applied for 5 days, after which the media was switched back to StemFlex media. Puromycin-resistant clones became visible after 7 days. Clones were pooled together, expanded, cryopreserved and later sorted into 96-well plates to ensure single clone formation by seeding one cell per well. The sorting was performed on a BD Aria II (Stanford Shared FACS Facility). The clone used in this study was selected based on its morphology and uniform expression of eGFP. Validation of hiPS genome integrity was performed by high density SNP arrays.

### Generation of hCOs and hStrOs from hiPS cells

hCOs and hStrOs were generated as previously described^18,24^. hiPS cells were cultured on plates coated with vitronectin (5 μg ml^−1^; Thermo Fisher Scientific, A14700) in Essential 8 medium. Cells were passaged every 4–5 d with UltraPure 0.5 mM EDTA (Thermo Fisher Scientific, 15575020). For the generation of 3D neural organoids, hiPS cells were incubated with Accutase (Innovative Cell Technologies, AT104) at 37 °C for 7 min and dissociated into single cells. One to two days before organoid formation, hiPS cells were exposed to 1% DMSO (Sigma-Aldrich, 472301) in Essential 8 medium. For aggregation into organoids, approximately 3 × 10^6^ single cells were added per well in AggreWell 800 plates in Essential 8 medium supplemented with the ROCK inhibitor Y27632 (10 μM), centrifuged at 100g for 3 min and then incubated at 37 °C with 5% CO_2_. After 24 h, organoids consisting of approximately 10,000 cells were collected from each microwell by pipetting the medium up and down in the well with a cut P1000 pipette tip and transferred into ultra-low attachment dishes (Corning, 3262) in Essential 6 medium (Thermo Fisher Scientific, A1516401) supplemented with the SMAD pathway inhibitors dorsomorphin (2.5 μM; Sigma-Aldrich, P5499) and SB-431542 (10 μM; R&D Systems, 1614). For the first 5 d, Essential 6 medium supplemented with dorsomorphin and SB-431542 was changed every day. Optionally, the Wnt pathway inhibitor XAV-939 (1.25 μM; Tocris, 3748) can be added with the two SMAD pathway inhibitors.

To generate hCOs, day 6 neural organoids were transferred to neural medium containing Neurobasal-A medium (Thermo Fisher Scientific, 10888022), B-27 supplement minus vitamin A (Thermo Fisher Scientific, 12587010), GlutaMAX (1:100; Thermo Fisher Scientific, 35050079), and penicillin-streptomycin (10,000 U ml^−1^, 1:100; Thermo Fisher Scientific, 15140122). The neural medium was supplemented with 20 ng ml^−1^ epidermal growth factor (EGF; R&D Systems, 236-EG) and 20 ng ml^−1^ basic fibroblast growth factor (FGF2; R&D Systems, 233-FB) for 16 d, with medium changed daily in the first 10 d and every other day for the following 6 d. From day 22, the neural medium was supplemented with brain-derived neurotrophic factor (BDNF; 20 ng ml^−1^; PeproTech, 450-02) and NT-3 (20 ng ml^−1^; PeproTech, 450-03), with medium changes every other day. From day 46, neural medium containing B-27 plus supplement (Thermo Fisher Scientific, A3582801) was used for medium changes every 4 d.

To generate hStrOs, day 6 neural organoids were transferred to neural medium containing Neurobasal-A medium, B-27 supplement minus vitamin A, GlutaMAX, penicillin-streptomycin, and supplemented with WNT pathway inhibitor IWP-2 (2.5 μM; STEMCELL Technologies, 72124) and recombinant Human/Murine/Rat Activin A (50 ng ml^−1^; PeproTech, 120-14P). From day 12, neural medium was additionally supplemented with retinoid X receptor agonist SR 11237 (100 nM; Tocris, 3411). From day 22, neural medium was supplemented with BDNF, NT-3, L-ascorbic acid 2-phosphate trisodium salt (AA; 200 μM; FUJIFILM Wako Chemical Corporation, 323-44822), N^6^,2’-O-dibutyryladenosine 3’,5’-cyclic monophosphate sodium salt (cAMP; 50 μM; MilliporeSigma, D0627) and cis-4,7,10,13,16,19-docosahexaenoic acid (DHA; 10 μM; MilliporeSigma, D2534). From day 42, neural medium was additionally supplemented with DAPT (2.5 μM; STEMCELL Technologies, 72082). From day 46, neural medium containing B-27 plus supplement was used for medium changes every 4 d.

### Real-time qPCR (RT-qPCR)

For RT-qPCR analysis, 3–4 hCOs were pooled per sample. mRNA from day 25 hCOs was isolated using the RNeasy Mini Kit (Qiagen, 74106). Template cDNA was prepared by reverse transcription using the SuperScript III First-Strand Synthesis SuperMix for qRT-PCR (Thermo Fisher Scientific, 11752250). RT-qPCR was performed using the SYBR Green PCR Master Mix (Thermo Fisher Scientific, 4312704) on a QuantStudio 6 Flex Real-Time PCR System (Thermo Fisher Scientific, 4485689). Primers used in this study are listed in **Supplementary Table 1**.

### Viral labeling

Viral infection of neural organoids was performed as previously described^24^. Briefly, neural organoids were transferred to a 1.5 ml Eppendorf tube containing 200 μl neural medium and incubated with the virus at 37 °C overnight. The next day, neural organoids were transferred to the fresh neural medium in ultra-low attachment plates (Corning, 3473).

The viruses used in this study are AAV-DJ-hSyn1::eYFP (Stanford University Neuroscience Institute Gene Vector and Virus Core, GVVC-AAV-16, 6597), AAV-DJ-hSyn1::mCherry (Stanford University Neuroscience Institute Gene Vector and Virus Core, GVVC-AAV-17, 5297), and AAV1-Syn::ChrimsonR-tdT (Addgene, 59171-AAV1)^22^.

### Integration of neural organoids with KiriE

The assembled KiriE on the culture chamber housed in the 10 cm tissue culture dish was brought into the biosafety hood. 70% IPA was perfused into the culture chamber and added to the 10 cm tissue culture dish to sterilize the whole setup. Then the 70% IPA was completely replaced with DPBS. The culture chamber was filled with laminin solution (1:100 dilution in DPBS; Sigma-Aldrich, L2020) and incubated overnight to promote the adhesion of KiriE.

The laminin solution was completely replaced by neural media before plating neural organoids on the KiriE. To ensure organoids are placed on the electrodes of KiriE, a homemade real-time imaging setup including a CMOS camera (OMAX) and a zoom lens was used during organoid integration. Media exchange was performed using a syringe pump media perfusion system (**Extended Data Fig. 5a**) starting from 2 days after integration.

### Reusability of KiriE

The cleaning procedure for the KiriE was adapted from a previously established protocol^34^. To dissolve the organoids that have been integrated with KiriE, the KiriE in the 3D culture chamber was incubated in 1× TrypLE Express (Thermo Fisher Scientific, 12604013) at 37 °C for 30 min and subsequently incubated in sodium dodecyl sulfate (Fisher Scientific, BP166500, diluted in water at a ratio of 4:100) overnight. The organoid residues were removed from the chamber with DPBS. To clean debris from the KiriE surface, a tablet of Ultrazyme enzymatic cleaner (Ultrazyme, 90278) was dissolved 15 ml of DPBS and incubated in the chamber at 37 °C overnight. 70% IPA was used to sterilize the culture chamber for the next culture.

### Generation and integration of cortico-striatal assembloids with KiriE

To simultaneously generate cortico-striatal assembloids and integrate them with KiriE, hCOs and hStrOs were differentiated separately and placed on the KiriE sequentially in close proximity. First an hStrO was placed on the electrodes in the center of KiriE, and two days later, an hCO was placed on the KiriE adjacent to the hStrO.

### Plating hCOs on the adhesive substrate

For plating hCOs on the adhesive substrate, 24-well glass-bottom plates (Cellvis, P24-0-N) were coated with Matrigel^14^. Specifically, the glass-bottom plates were coated with a 2% Matrigel solution (Corning, 356234) in DPBS overnight. On the following day, the Matrigel solution was removed and a single hCO was placed in the center of each well. Neural medium was added and changed according to the same schedule as typical hCO cultures. The plates were incubated at 37 °C with 5% CO_2_.

### Live-cell imaging and analysis

The multifunctional culture chamber allows for live-cell confocal imaging of neural organoids, assembloids, and KiriE through the glass bottom. The culture chamber was incubated in an environmentally controlled chamber on a Leica Stellaris 5 inverted confocal microscope with a motorized stage for 15 min before imaging.

The cell viability assay was conducted using SYTOX™ deep red nucleic acid stain for dead cells (Thermo Fisher Scientific, S11381), as per the manufacturer’s instructions. In brief, LMNB1-mEGFP hCOs were incubated with SYTOX, which was diluted in neural medium to a 1:1,000 ratio, in a CO_2_ incubator at 37 °C for 15 min. Subsequently, the hCOs were rinsed twice with neural medium before undergoing live-cell confocal imaging.

Imaging data was analyzed as previously described^20^. Briefly, Imaris 9 software (Bitplane) was used for visualization and segmentation. A combination of Imaris, ImageJ (National Institutes of Health) and MATLAB (MathWorks) was used for quantitative analysis of fluorescence intensity as a function of the distance from the KiriE. Imaris was used to define the KiriE structures and the 3D volume of hCOs. ImageJ was used to export fluorescence intensities, distances from the KiriE and associated 3D coordinates. MATLAB was used to calculate fluorescence intensity as a function of the distance from the KiriE. The average fluorescence intensity values for all voxels with distances binned over an interval of 10 μm were normalized against the baseline value as the average fluorescence intensity of all voxels 150**–**160 μm away from KiriE. The cell counting for the SYTOX cell viability assay was conducted using Imaris 9 software. Due to the far-red background, only nuclei with clear, round morphology were quantified as SYTOX^+^ cells. Two individuals carried out the cell counting independently, and the mean values were reported as the final results.

Characterization of size and circularity of hCOs on KiriE was performed using ImageJ.

### FIB-SEM imaging

FIB-SEM samples were prepared using our previously developed ultra-thin plasticization protocol^35,36^. The buffer is 0.1 M sodium cacodylate (Electron Microscopy Sciences) unless stated otherwise. Briefly, specimens were fixed overnight in 2.5 % glutaraldehyde (Electron Microscopy Sciences) at 4 °C, washed 3 times in chilled buffer, and quenched with 20 mM glycine (Merck & Co.). Each organoid was post-fixed in 2% osmium tetroxide (Electron Microscopy Sciences) and 1% potassium ferrocyanide (Electron Microscopy Sciences), washed 3 times in chilled buffer, then incubated with 1% thiocarbohydrazide (Electron Microscopy Sciences) in distilled water. After 3 washes in water, samples were incubated again in 1% osmium tetroxide in water for 1 h, and then rinsed in water. Subsequently, the en bloc staining was carried out overnight with 1% uranyl acetate (Electron Microscopy Sciences) at 4 °C followed by 3 washes in water. Prior to dehydration, 0.15 % tannic acid aqueous solution was added. Dehydration was achieved by immersing the sample in an increased concentration of ethanol (Merck & Co.) starting from 30% to 100%. For resin embedding, a mixture of low viscosity Spurr’s resin (Electron Microscopy Sciences) in ethanol was added with the following ratio: 1:3 (2 h); 1:2 (2 h); 1:1 (overnight); 2:1 (2 h); 3:1 (2 h). Then, specimens were embedded in fresh resin for 24 h and then placed in vertical position for 2**–**3 h to remove the excess resin. Each sample was washed for 3**–**5 seconds with absolute ethanol to further remove excess resin on the surface. Polymerization was initiated by moving the sample into an oven at 70 °C. After 24 h polymerization, samples were cut into smaller parts for easy mounting and imaging. Each organoid part was mounted on a 12 mm aluminum stub using silver paste. Mounted samples were coated with 20 nm of gold by using HR 208 Sputter coater (Cressington), then loaded into FIB-SEM chamber (Helios CX5, Thermo Scientific). The sample surface was first scanned with a 3 kV electron beam to identify the region of interest (ROI). An ROI for regular cross section was located for milling with a nominal depth of 15**–**20 μm, length of 30**–**40 μm and the width varied in a range of 30**–**100 μm. After polishing the region of interest, image acquisition was performed in immersion mode, with a TLD detector with back scattered configuration at 2**–** 3 kV, 0.17**–**0.34 nA of current in a range of magnification from 5kx to 40kx.

### Cryopreservation and immunohistochemistry

Cryopreservation and immunohistochemistry were performed as previously described^24,33^. In brief, neural organoids were fixed in 4% paraformaldehyde (PFA; Electron Microscopy Sciences, 15710) in PBS at 4 °C for 2 h. They were washed in PBS and transferred to 30% sucrose in PBS for 2 d until they sank in the solution. Then the neural organoids were rinsed and embedded in a mixture comprising optimal cutting temperature (OCT) compound (Tissue-Tek OCT Compound 4583, Sakura Finetek) and 30% sucrose in PBS at a 1:1 ratio. 20–30-μm-thick sections were cut using a Leica Cryostat (Leica, CM1860). Cryosections were washed with PBS to remove excess OCT and blocked in 10% normal donkey serum (NDS; Sigma-Aldrich, S30-M) and 0.3% Triton X-100 (Sigma-Aldrich, T9284) at RT for 1 h. Then the sections were incubated with primary antibodies diluted in PBS containing 2% NDS and 0.3% Triton X-100 at 4 °C overnight. Next day, excess primary antibodies were washed away by PBS, and the cryosections were incubated with secondary antibodies in PBS containing 2% NDS and 0.3% Triton X-100 at RT for 1 h. Following PBS washes, nuclei were counterstained with Hoechst 33258 (1:10,000 dilution; Thermo Fisher Scientific, H3569, 2086723). Finally, slides were mounted with coverslips using Aqua-Poly/Mount (Polysciences, 18606). The slides were kept in the dark for at least 24 h before microscopic imaging.

The following primary antibodies were used for staining: anti-SOX2 (rabbit, Cell Signaling Technology, 3579S, 8, 1:300 dilution), anti-MAP2 (guinea pig, Synaptic Systems, 188004, 3-37, 1:200 dilution), anti-GFAP (rabbit, Dako, Z0334, 41259205, 1:500 dilution), cleaved caspase-3 (Asp175) antibody (rabbit, Cell Signaling Technology, 9661, 45 and 47, 1:200 dilution), anti-neurofilament heavy polypeptide antibody (mouse, Abcam, ab7795, GR299862-23, 1:200 dilution), anti-TBR1 (rabbit, Abcam, ab31940, GR3215726-2, 1:100 dilution), anti-SATB2 (mouse, Abcam, ab51502, GR3235758-6, 1:50 dilution). The following secondary antibodies were used at a 1:1,000 dilution for staining: Alexa Fluor 647 AffiniPure donkey anti-rabbit IgG (H+L) (Jackson ImmunoResearch, 711-605-152), Alexa Fluor 647 AffiniPure donkey anti-guinea pig IgG (H+L) (Jackson ImmunoResearch, 706-605-148), donkey anti-rabbit IgG (H&L) highly cross-adsorbed secondary antibody Alexa Fluor 488 (Thermo Fisher Scientific, A-21206), donkey anti-mouse IgG (H+L) highly cross-adsorbed secondary antibody Alexa Fluor 647 (Thermo Fisher Scientific, A-31571), donkey anti-mouse IgG (H+L) highly cross-adsorbed secondary antibody Alexa Fluor 488 (Thermo Fisher Scientific, A-21202).

The average fluorescence intensity values for all voxels with distances binned over an interval of 1 μm were normalized against the baseline value as the average fluorescence intensity of all voxels 50-51 μm away from KiriE.

### Single cell RNA sequencing and data analysis

Dissociation of hCOs into single cells for scRNA-seq was performed as previously described^37^ with minor optimizations. 3–4 hCOs at day 100 were pooled from KiriE-integrated condition or control condition from each hiPS cell line. hCOs were transferred to a 6-well plate (Corning, 3506) and incubated for 45–60 min at 37 °C with 3 ml enzymatic dissociation solution. The enzymatic solution consisted of 30 U/ml papain (Worthington Biochemical, LS003127), 1x EBSS (Sigma-Aldrich, E7150), 0.46% D(+)-glucose, 0.5 mM EDTA, 26 mM NaHCO_3_, 10 μM Y-27632, 125 U/ml

Deoxyribonuclease I (Worthington Biochemical, LS002007), and 6.1 mM L-cysteine (Sigma-Aldrich, C7880). Upon the completion of papain incubation, samples were collected in a 15 ml Falcon tube and centrifuged at 1,200 rpm for 1 min. After removing the supernatant, the samples were washed with 1 ml inhibitor solution with 2% trypsin inhibitor (Worthington Biochemical, LS00308) and resuspended in 1 ml of the same solution for trituration. Upon the completion of trituration, 1 ml inhibitor solution with 4% trypsin inhibitor was slowly added beneath the cell suspension to create a gradient layer. The gradient solution was centrifuged at 1,200 rpm for 5 min. Cell pellets were resuspended in 0.2% BSA/PBS and passed through a 40 μm cell strainer (Corning, 352340). The inhibitor solution differs from the enzyme solution in that it does not contain papain nor EDTA. All centrifugation steps were performed at RT.

Approximately 8,000 cells were loaded onto a Chromium Next GEM Single Cell 3’ GEM Kit v3.1 chip (10x Genomics, PN-1000123). Libraries were generated with the Chromium Next GEM Single Cell 3’ Library Kit v3.1 (10x Genomics, PN-1000157) using the Dual Index Kit TT Set A (10x Genomics, PN-1000215). Libraries were sequenced and demultiplexed by Admera Health using the Illumina NovaSeq S4 2 × 150. Mapping to the human reference genome (Human (GRCh38) 2020-A provided by 10x Genomics) and generation of count matrices (including introns) were performed using 10x

Genomics Cloud Analysis using Cell Ranger software (version 7.0.0). Samples were aggregated including normalization for mapped reads using the aggregate function in 10x Genomics Cloud Analysis. Downstream analyses were performed on the filtered count matrices in RStudio (version 1.4.1717 and R version 4.1.1) using the Seurat R package (version 4.1.1)^38^. Cells with more than 9,000 or less than 1,000 detected genes or with mitochondrial content higher than 10% were excluded. Genes that were not expressed in at least three cells were not included in the analysis. Gene expression was normalized using a global-scaling normalization method (normalization method, ‘LogNormalize’; scale factor, 10,000), and the 2,000 most variable genes were selected (selection method, ‘vst’) and scaled (mean = 0 and variance = 1 for each gene) before principal component analysis. The top 21 principal components were used for clustering (resolution of 0.6) using the ‘FindNeighbors’ and ‘FindClusters’ functions, and for visualization with UMAP. Clusters were grouped based on the expression of known marker genes and differentially expressed genes, and more resolved clusters and the top ten genes are shown in **Extended Data Fig. 4e,f**. Then the standard workflows for visualization with UMAP and clustering were used.

### Electrophysiological recording, optogenetic and pharmacological modulation, and data analysis

The electrophysiology of neural organoids and assembloids was recorded by KiriE in the multifunctional culture chamber in the cell culture incubator. The FFC was connected to an Intan RHS 32-channel stim/recording headstage (Intan Technologies) through a homemade printed circuit board. Stainless steel wires were used as references. Electrophysiological recordings were acquired at a 20 kHz sampling rate. The electrical impedance of each electrode at 1 kHz was also measured. All working channels are included in the reported data. The determination of working channels is based on impedance measurements with an upper cutoff value of ∼0.8 MΩ and a lower cutoff value of ∼0.1 MΩ.

For optogenetic stimulation, AAV1-Syn::ChrimsonR-tdTomato^+^ cells in hCOs or in hCOs of cortico-striatal assembloids were activated with 590 nm light through a fiber optic cannula (Thorlabs) from the bottom of the culture chamber (**Extended Data Fig. 8k**). A custom-designed 3D printed adaptor was used to attach the fiber optic cannula vertically to the shelf in the incubator. The optogenetic stimulation is a pulse train of 1 s light pulse every 10 s. The pulse was generated by a Cyclops LED driver (Open Ephys), which was controlled by Arduino Uno (Arduino).

The application of pharmacology was performed using the built-in perfusion system of the multifunctional culture chamber. Upon completion of measurement, the drugs were completely washed out by perfusing excessive neural medium through the culture platform. Firing rates of individual channels were compared before and after treatment with potassium channel blocker 4-AP (50 μM; Tocris, 0940), sodium channel blocker TTX (1 μM; Tocris, 1069), AMPA receptor antagonist NBQX (20 μM, Tocris, 0373) and NMDA receptor antagonist APV (50 μM, Tocris, 0106).

Acute electrophysiological recordings using silicon shank electrodes were conducted following our recently published protocol^39^. Briefly, the neural organoids were embedded in 3% low-melting agarose (IBI Scientific, IB70056) and then transferred into artificial cerebrospinal fluid (aCSF) containing 124 mM NaCl, 3 mM KCl, 1.25 mM NaH_2_PO_4_, 1.2 mM MgSO_4_, 1.5 mM CaCl_2_, 26 mM NaHCO_3_ and 10 mM D-(+)-glucose with the addition of GlutaMAX (Gibco, 35050061). The organoids were perfused with aCSF maintained at 37 °C. Electrophysiological recordings were acquired using an acute 32-channel probe (Cambridge NeuroTech) connected to the Intan 1024-channel recording controller (Intan Technologies).

Data analysis was performed in Python. The recordings generated by the Intan RHS interface are stored in an .rhs format which can be opened via the *pyintan* package in Python. The recordings were made into one .npy (*numpy*) format. The recordings were filtered with a 3^rd^ order Butterworth bandpass filter between 150 Hz and 9000 Hz. The filter was applied in a non-causal way to preserve the shape of action potentials and to help differentiate artifacts from real electrophysiological signals. This filtering is performed with the *scipy.signal* package using its functions *butter* and *filtfilt*.

Bandpass-filtered large deviations from baseline were collected via *scipy.signal’*s *find_peaks* function. The standard deviation was computed as a sliding standard deviation in one-second windows using Fourier-space convolutions. First, all signals deviating more than 5 standard deviations from the baseline (either positive or negative) were collected by taking 1.5 milliseconds before and after their maximal point (minimal if negative) for a total of 3 ms as depicted in **Extended Data Fig. 8b**. The detected signals from all electrodes were grouped for a given recording and clustered according to their shape (**Extended Data Fig. 8c,d**). Two-component UMAP (*umap-learn* package) dimensionality reduction was followed by a density-based clustering (DBSCAN function of the *sklearn.cluster* package). The identified clusters correspond to spikes of similar shape (**Extended Data Fig. 8e**). Representative spikes are presented by averaging all the signals belonging to a given cluster. Spikes representing real signals (but not artifacts) were extracted from the recordings by means of z-normalized template-matching (*match* function of the *stumpy* package; see **Extended Data Fig. 8f**). Overall, this method provides a noise-robust, shape-based spike detection in our electrophysiology recordings.

After spike detection, a binary array with the same length and number of channels as the recording was built. This array contains 1s at the timestamps where spikes were detected and 0s everywhere else. A gaussian kernel of 3 s in length and 0.5 s of standard deviation was convolved over the binary array for every channel. This convolution operation with such a kernel behaves as a smooth, sliding event counter. The resulting array were displayed in false color to convey the instantaneous firing rate of the neural organoids.

## Statistics

Comparisons between experimental groups were made using two-tailed *t*-tests or one-way analysis of variance (ANOVA) tests with multiple comparisons. *P* values of less than 0.05 were considered statistically significant.

## Data Availability

Single cell gene expression data are available under the Gene Expression Omnibus accession number GSE221032 (token kzsbqacydtwllef). The data that support the findings of this study are available from the corresponding authors upon request.

## Code Availability

The code used for data analysis is available from the corresponding authors upon request.

## Author Contributions

B.C., S.P.P., X.Y., and C.F. conceived and designed the project. C.F. developed the Python code for scripted pattern design, performed the FEM simulations, and optimized the KiriE parameters. X.Y. and T.L.L. maintained and differentiated hiPS cells into hCOs. Y.M. maintained and differentiated hiPS cells into hStrOs. C.F., C.T.T., X.Y., and T.J.Z. fabricated the devices. X.Y., B.C., C.F., and T.L.L. designed the culture chamber and X.Y. characterized the media exchange system. X.Y., C.F., and T.L.L. developed the device assembly protocol and X.Y. developed direct contact interfacing protocol. T.J.Z., X.Y., T.L.L., and C.F. assembled the devices. X.Y. and T.L.L. integrated neural organoids with devices and maintained the cultures. X.Y., S.K., and X.C. performed scRNA-seq experiments. X.Y., S.K., and Y.M. analyzed scRNA-seq data. X.Y. and J.P.M. performed qPCR experiments. J.P.M. characterized the morphology of hCOs. X.C. generated the CAG::eGFP hiPS cell line. X.Y. performed immunocytochemistry, confocal imaging, and data analysis. X.Y. and T.L.L. performed electrophysiological recordings. V.M., F.S., and C.T.T. performed electron microscopy imaging. C.F. wrote the code and analyzed the electrophysiological data. X.Y., C.F., B.C., and S.P.P. wrote the manuscript with input from all authors. B.C. and S.P.P. supervised the work.

## Acknowledgements

We thank members of the Paşca and Cui labs for helpful discussions and technical support. In particular, we thank S.J. Yoon for sharing hCOs for initial testing, Y. Yang for help with initial testing, J.I. Kim, M.Y. Li and O. Revah for discussions on electrophysiological recordings.

This work was financially supported by the National Institutes of Health (R01NS121934, R01HL165491, R35GM141598) (to B.C.), the Stanford Big Idea Project on Brain Organogenesis (Wu Tsai Neurosciences Institute) (to S.P.P. and B.C.), the David and Lucile Packard Foundation (to B.C.), the Kwan Funds (to S.P.P.), the Senkut Funds (to S.P.P.), the Coates Foundation (to S.P.P.), the Ludwig Family Foundation (to S.P.P.), the Alfred E. Mann Foundation (to S.P.P.), the Marie Curie Fellowship (to C.F.), and an EMBO Postdoctoral Fellowship (ALTF 321-2021) (to S.K.). S.P.P. is a New York Stem Cell Foundation (NYSCF) Robertson Stem Cell Investigator, a CZ BioHub Investigator and a CZI Ben Barres Investigator. Part of this work was performed at the Stanford Nano Shared Facilities (SNSF), supported by the National Science Foundation under award ECCS-2026822.

## Competing Interests

Stanford University holds patents for the generation of cortical organoids/spheroids (listing S.P.P. as an inventor). Stanford University has filed a provisional patent application that covers the devices, systems, and methods of KiriE (listing B.C., S.P.P., X.Y., and C.F. as inventors).

**Extended Data Fig. 1.**
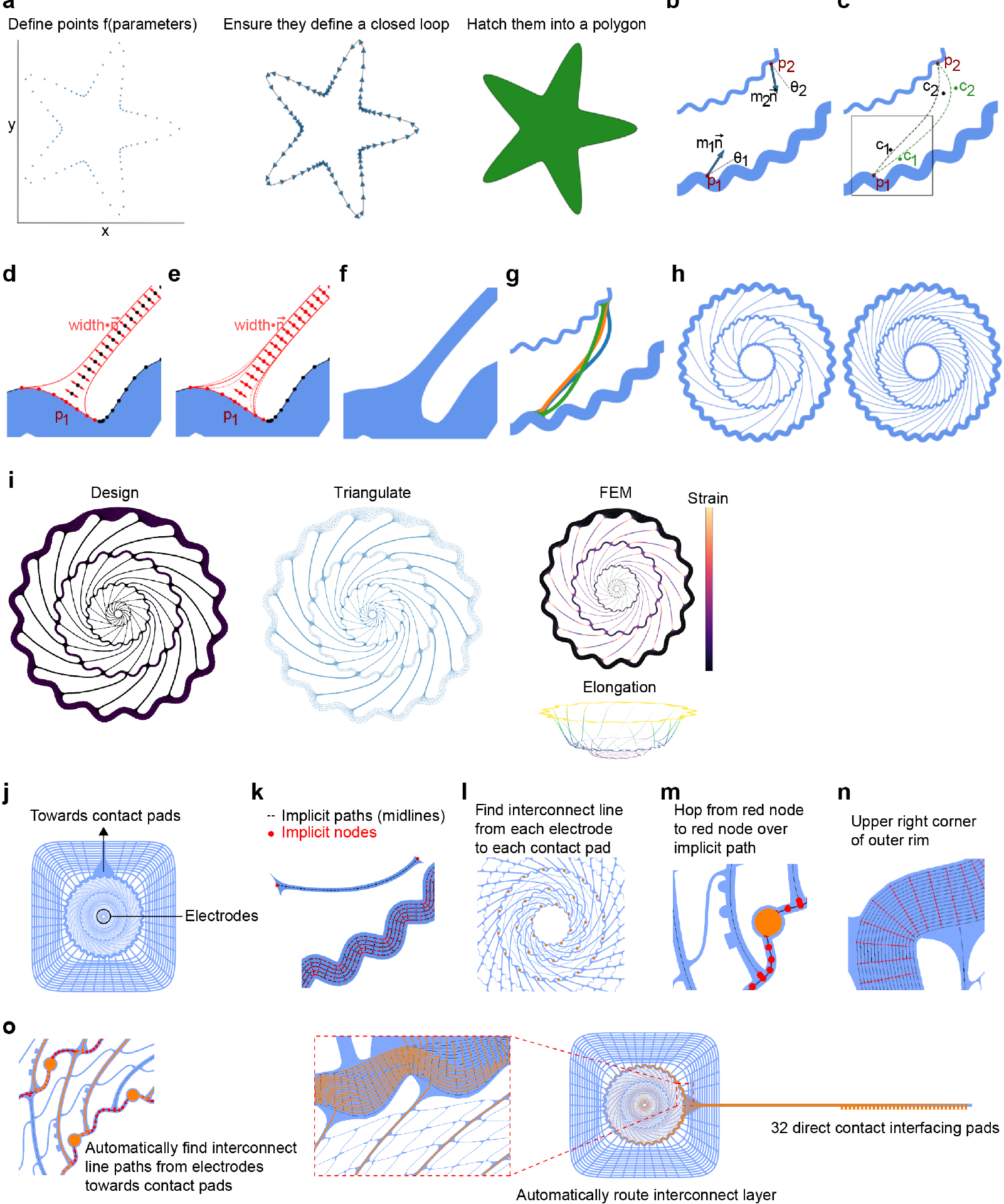
Script-assisted design to generate KiriE patterns and automated electrode routing. **a**, The procedure that generates polygon patterns for photolithography. The first step is to generate a list of (x,y) coordinates. The second step is to connect these coordinates into a continuous, directed closed loop. Finally, these closed loops are filled into polygons utilizing a Python geometry library, *PHIDL*. The infill decides whether the lithography mask is transparent or opaque and the output GDS-II file is compatible with photolithography. **b**, The latch design. The starting points (p), the normal vectors at those points 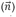 and optionally a rotation of these normals (θ) serve as input parameters to build a latch connecting two polygons. The magnitude (m) of either normal vector is adjustable. **c**, The starting points and the normal vectors (rotated normal in green) define 4 points p1, p2, c1, c2. These points are control points for a NURBS curve which defines the center of the latch. **d**, The normal vectors of the newly generated curve are used to give it a width. The end of the latch is smoothly snapped onto the polygon to make sure that the contact is flush (no holes or anomalies for lithography). **e**, After the latch is constructed, the connection is adjusted to smooth out any sharp joins at the attachment (see **Fig. 1e**). **f**, The resulting latch is integrated into the design. **g**, A variety of latches can be produced by simply changing the input parameters to the script. **h**, The spiral KiriE is programmatically generated by inputting the number of latches, rings, size, latch parameters, etc. Complex KiriE patterns are generated and iteratively modified. **i**, The simulation workflow. The design file is turned into a triangulated mesh by *pygmsh*. This allows mechanical simulations with FEM to calculate KiriE vertical elongation and strain distribution. **j**, The layout of spiral KiriE pattern that shows the device area (inside the outer ring) and the mesh outside the device area for attaching to the supporting structure. **k**, To automatically route the electrode lines, implicit nodes (red circles) and implicit paths (black dotted line) were generated using the Polygon class in *PHIDL*. The implicit paths were used to construct the metal line routes. The nodes were used for path-finding through the device. **l**, Each electrode (yellow) is routed through an individual non-intersecting metal line towards the contact pad. **m**, For automated path finding, the implicit nodes (red) define a graph and the nodes of distinct polygons are connected in the form of an edge inside the graph if they are close enough. **n**, Inset of the outer wavy ring of the device showing the implicit nodes and paths inside the polygon. **o**, All the electrodes’ nodes are connected to a common “virtual” drain node, and the interfacing pads’ nodes to a “virtual” source node. The drain and source nodes serve the purpose of setting up an Edmond-Karps flow algorithm through the graph. A node-disjoint flow through the graph was found using the *Networkx* package, which translated to non-intersecting routes through the red nodes from electrode to pad. The implicit paths connected to red nodes were used to outline the exact shape of the metal line route (yellow paths). The right inset shows metal paths from electrodes to pads as they travel through the outer ring, towards the pads, without intersecting.

**Extended Data Fig. 2.**
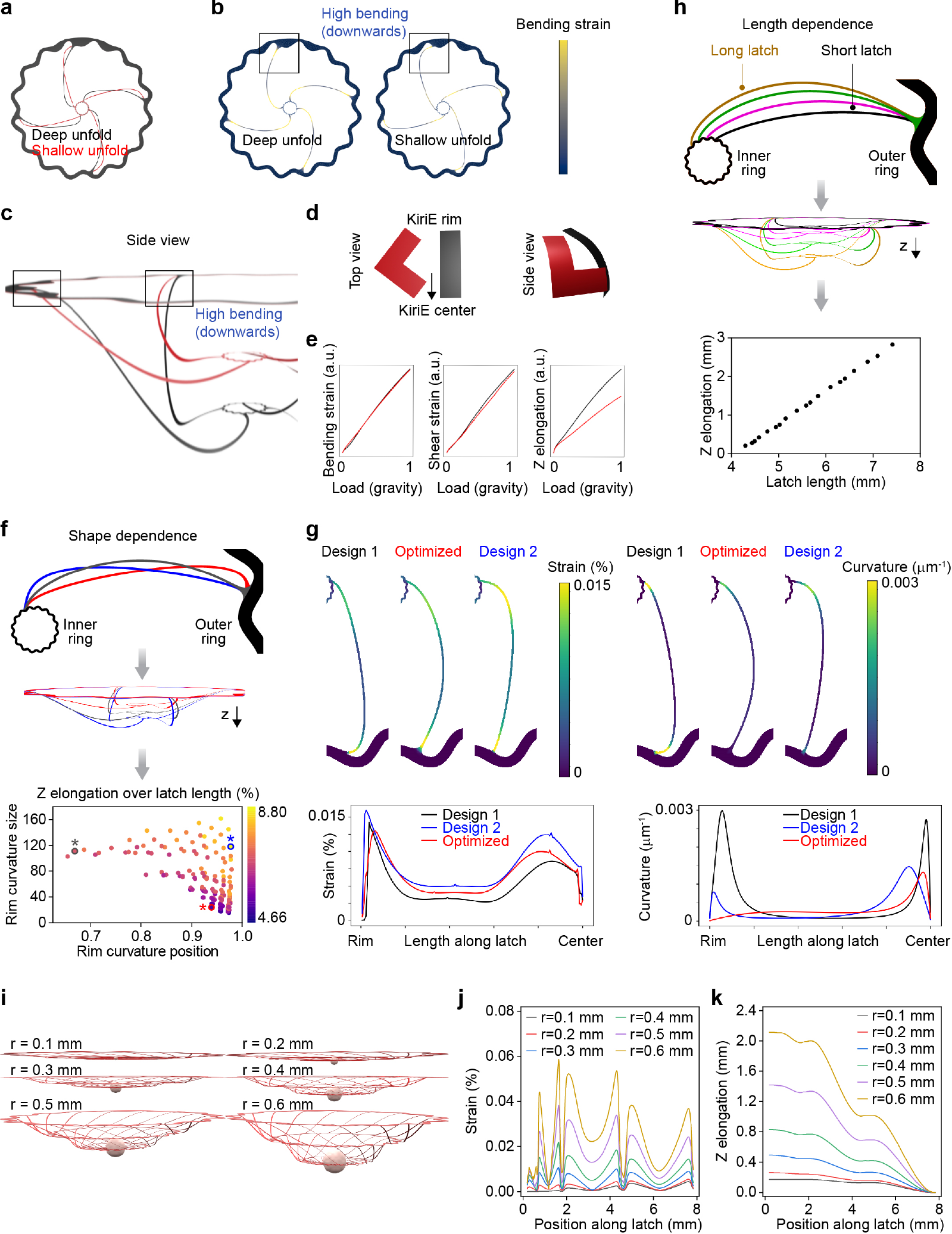
Influence of latch shape, latch length, and organoid size on vertical elongation in the spiral KiriE design. **a**, Two latch shapes of equal length connect two rings with identical connecting points. **b**, The bending strain that leads to the unfolding occurs at the outer rim of the design. The outer rim is fixed, while the rest of the design experiencing load is flexible. Therefore, the edge that is closest to the outer rim supports the most load and bends the most. **c**, A side view of the extended KiriE shows that both latches perform their downward bend close to the outer rim, exactly where the bending strain is highlighted as important in **b**. The red latch extends about half as deep as the black latch under equal load. **d**, Exaggerated representation of a red latch and a black latch for explanatory purposes. The red latch performs a sharp turn at the initial segment of the latch, represented here by a right angle. The red latch achieves less extension than the black latch is because the part of the latch that comes after the bent part does not proceed in the same direction, but veers off to a different direction. In the case of the black latch, the initial segment bends and points downwards and the rest of the latch follows that direction in a nearly straight direction. In contrast, the part of the red latch that comes after the right angle does not point to the direction that would achieve more extension. **e**, The achieved extension depends on geometrical arguments. The bending and shear strains for black and red latches are nearly identical, suggesting that it is not a simulation locking issue that prevents the design from extending further. **f**, FEM simulation shows that the vertical deformation is dependent on latch shape when the latch length and the connecting points are fixed. The three data points with asterisks correspond to the three latch shapes. **g**, The three highlighted latch shapes from **f** are reproduced here to highlight their difference. While design 2 latch vertically extends ∼15% more than the optimized latch, it has a higher curve near the rim, which increases local strain accumulation. The optimized shape is designed according to **Extended Data Fig. 1j–o** (**Methods**) such that for a given length, the total curvature is minimized. This produces a smooth latch which has sufficient (not maximal) vertical extension but has a low maximal strain along the latch. **h**, FEM simulation shows that the vertical deformation is dependent on the latch length, which is determined by the location of the connection on the inner ring. **i**, Schematic of KiriE vertically deformed by a neural organoid with a radius of 0.1, 0.2, 0.3, 0.4, 0.5, and 0.6 mm, respectively. **j**, The plot of strain along the metal interconnect of a microelectrode. **k**, The plot of Z elongation along the metal interconnect of a microelectrode.

**Extended Data Fig. 3.**
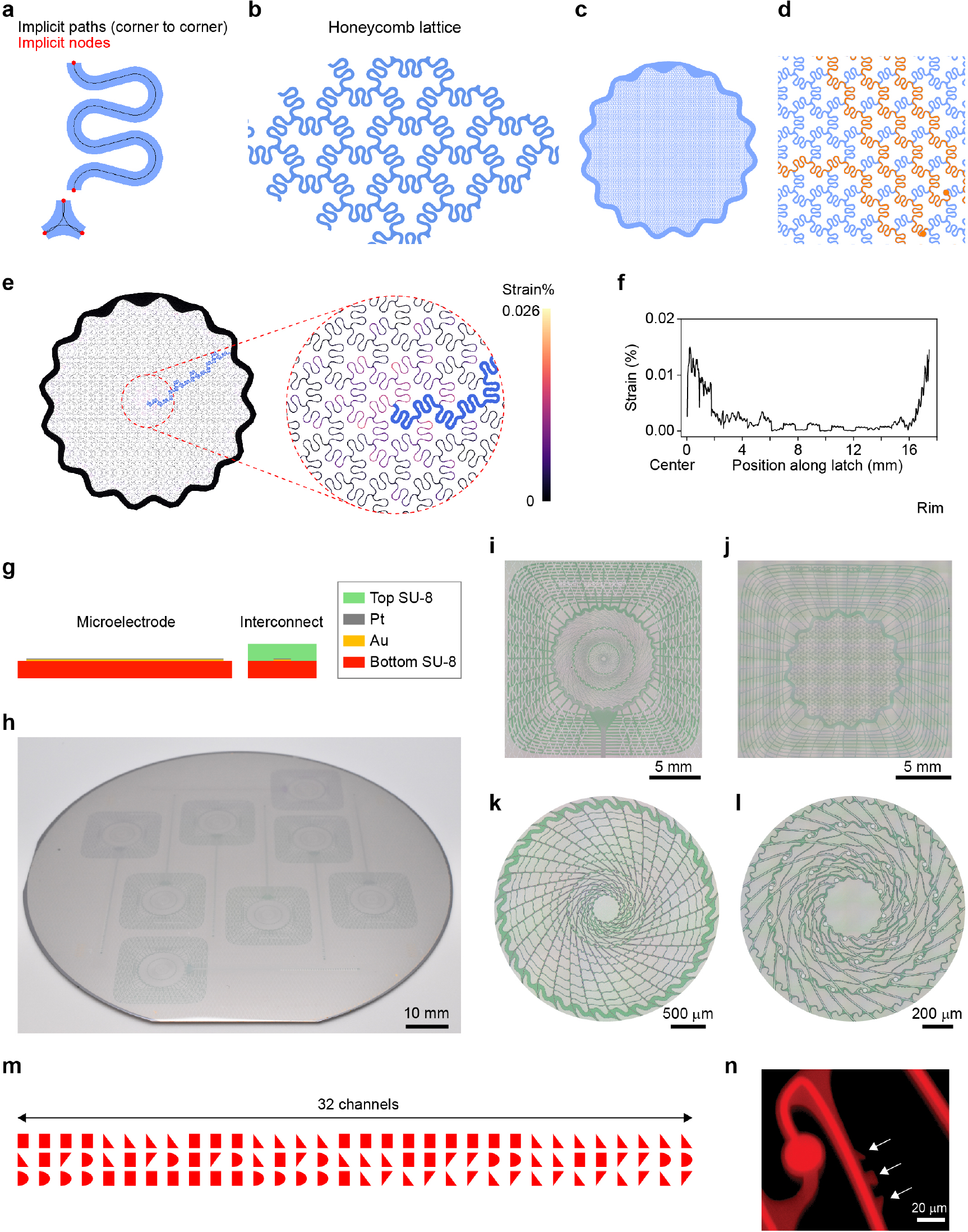
Honeycomb KiriE design and nanofabrication of KiriEs. **a**, A short serpentine line connected to a 3-way connector is the core of the honeycomb pattern. The algorithm for automated electrode routing is the same as spiral design as shown in **Extended Data Fig. 1j–o**. The only differences are locations of implicit nodes and paths for the basic polygons used to make the device. **b**, The tiling of the serpentine is done via the 3-way connector and tiled until it reaches a preset perimeter. **c**, The final honeycomb pattern with the motif tiled hexagonally within the confines of the outer ring (1 cm diameter). **d**, The yellow lines are the automatically routed electrode lines and the yellow circles are the measuring electrodes. **e**, Strain distribution in a simplified honeycomb KiriE under the weight of KiriE itself and a 1.2 mm-diameter organoid in medium. The strain is appreciable at the center because of the way the design gains vertical extension. For the honeycomb design, the serpentine loop stretches to gain vertical elongation, which strains at the central part of the serpentines. **f**, The strain along an interconnect line (highlighted in blue) is plotted from the center to the rim. The strain increases as the line gets closer to the center. **g**, Left: the cross-sectional schematic of the Pt/Au microelectrode (150 nm thick) supported on a layer of SU-8 (450 nm thick). Right: the cross-sectional schematic of an interconnecting electrode line that shows Pt/Au is sandwiched between two insulating layers of SU-8 (900 nm thick). **h**, A photograph showing 8 fabricated spiral KiriEs on a 4 inch (100 mm) silicon wafer before etching the Ni sacrificial layer. **i** and **j**, Optical images showing the entire pattern of (**i**) spiral KiriE design and (**j**) honeycomb KiriE design. The mesh lines outside the 1-cm wavy ring are designed to adhere the device to a PDMS top support. **k,l**, Zoomed-in optical images of the 32 electrodes located in the center of spiral KiriE. **m**, 32 channel-indexing barcodes incorporated into the KiriE design. **n**, A confocal image of an electrode with its associated channel-indexing barcode, a 3-digit geometrical shape (arrows).

**Extended Data Fig. 4.**
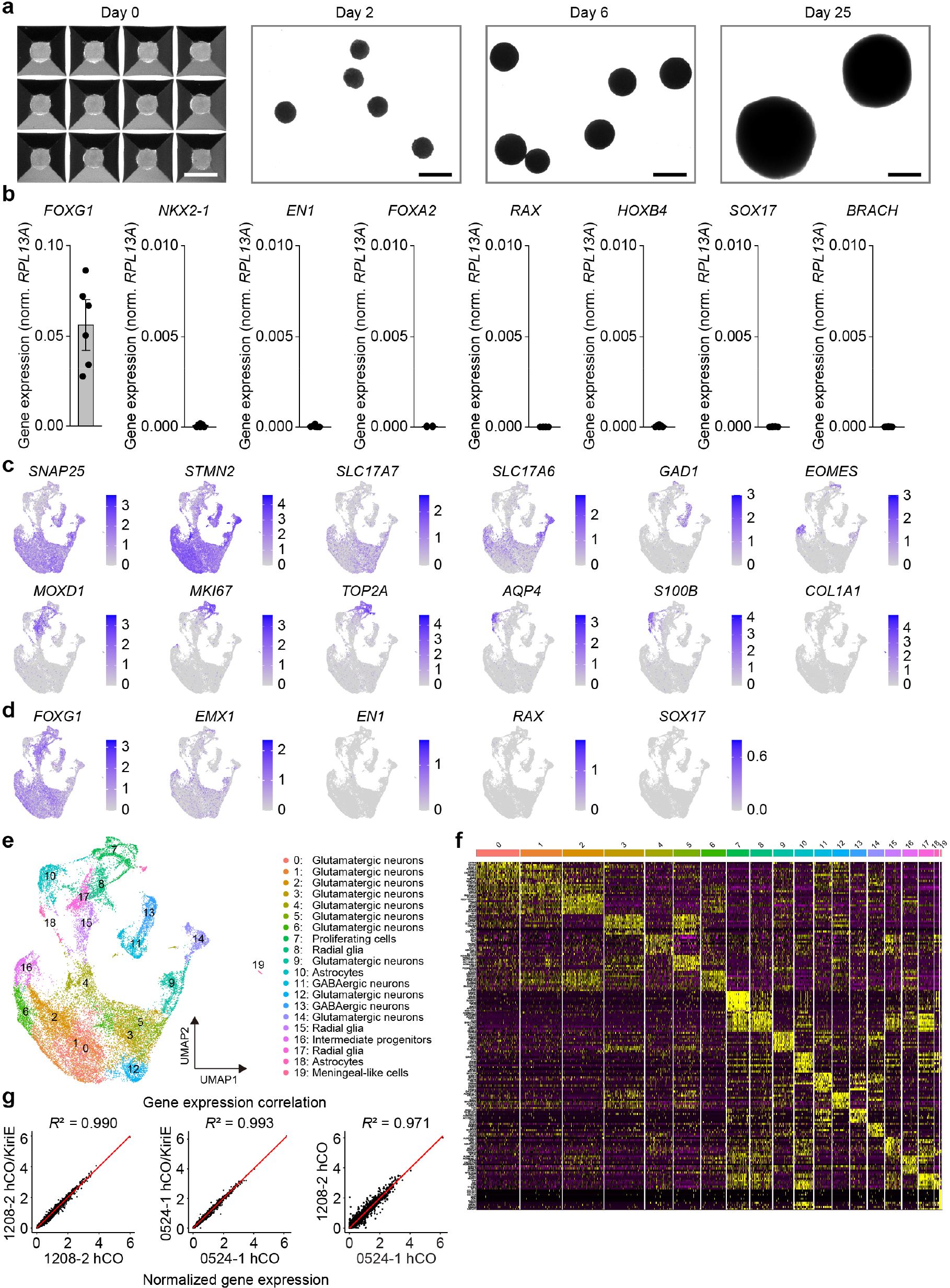
Generation and characterization of hCOs, and single cell transcriptomic characterization of KiriE-integrated hCOs. **a**, Representative hCOs generated in the AggreWell microwell plate at day 0, and hCOs cultured in ultra-low attachment plates at days 2, 6 and 25 of differentiation. Scale bars, 500 μm. **b**, Gene expression of *FOXG1, NKX2-1, EN1, FOXA2, RAX, HOXB4, SOX17*, and *BRACH* relative to housekeeping gene *RPL13A* in hCOs at day 25 of differentiation (*n* = 6 samples from three hiPS cell lines). Data are presented as mean ± s.e.m. **c**, UMAP visualization of expression of selected genes in the hCO scRNA-seq data at day 100 of differentiation (*n* = 25,546 cells from *n* = 4 samples of control hCOs and KiriE-integrated hCOs from two hiPS cell lines). **d**, UMAP visualization of expression of forebrain (*FOXG1*), dorsal forebrain (*EMX1*), midbrain (*EN1*), hypothalamus (*RAX*), endoderm (*SOX17*) markers in hCOs. **e**, UMAP visualization of the resolved scRNA-seq data of the aggregated data from control hCOs and KiriE-integrated hCOs. **f**, Heatmap of the top 10 genes in each cluster. **g**, Pearson correlations of the normalized average gene expression of 1208-2 hCO/KiriE vs. 1208-2 hCO (*r*^2^ = 0.990), 0524-1 hCO/KiriE vs. 0524-1 hCO (*r*^2^ = 0.993), and 1208-2 hCO vs. 0524-1 hCO.

**Extended Data Fig. 5.**
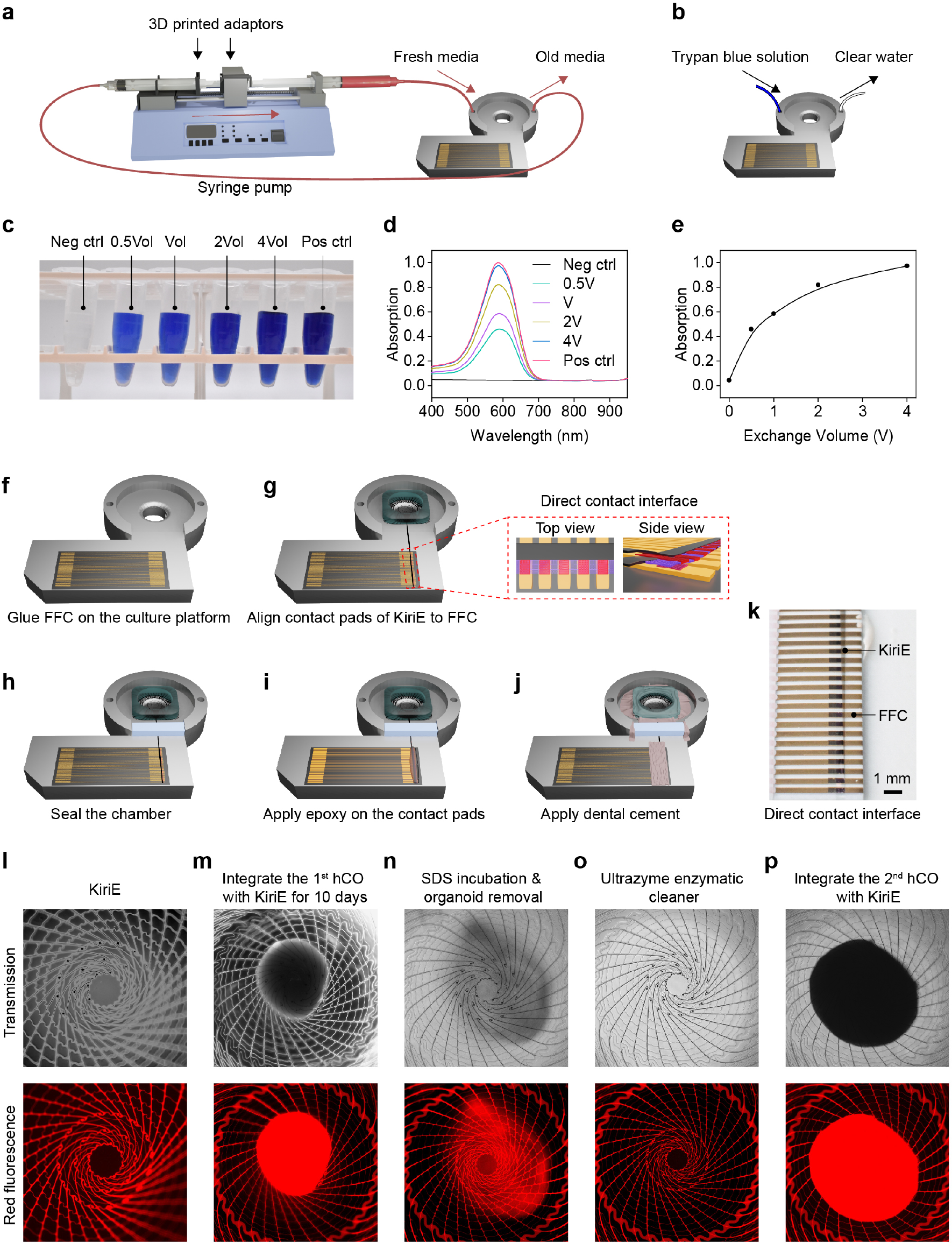
Platform for long-term KiriE-organoid integration and demonstration of KiriE reusability. **a**, Schematics of the KiriE-organoid culture and media exchange system. Inlet and outlet of the culture chamber are connected to two syringes that are attached to a syringe pump through custom-designed 3D printed adapters. **b**, An illustration of how the medium exchange efficiency is measured. **c**, Photograph of fluid collected from the culture chamber after replacing 0.5, 1, 2 and 4 times of the chamber volume of clear water with trypan blue solution, as well as negative (clear water) and positive (trypan blue solution) controls. **d**, UV-Vis absorption spectra of trypan blue in the fluid normalized against positive control. **e**, Maximum absorption as a function of media exchange volume. **f–j**, Illustrations of the KiriE device assembly process. **f**, Glue an FFC on the surface of the culture chamber. **g**, Transfer the KiriE device with a PDMS support onto the center of the culture chamber and align the contact pads of KiriE to FFC. Inset shows the top view and side view of the direct contact interface. **h**, Glue a 3D printed bar to the opening of the chamber with Metabond cement to seal the chamber. **i**, Aligned contact pads were fully dried by surgical spears (Braintree Scientific) and fixed in place using epoxy adhesive. **j**, The interface was covered with Metabond cement. **k**, An image of the contact pads on the KiriE tail aligned with the electrode pads of the FFC through direct contact interfacing. **l–p**, KiriE is reusable after a two-step cleaning. The transmission images and corresponding red fluorescence images of KiriE (**l**) before integration with an hCO, (**m**) after integration with the 1^st^ hCO, (**n**) after SDS treatment and organoid removal, (**o**) after Ultrazyme® enzymatic cleaner treatment, and finally (**p**) after integrating with the 2^nd^ hCO illustrate the complete cycle of KiriE reuse.

**Extended Data Fig. 6.**
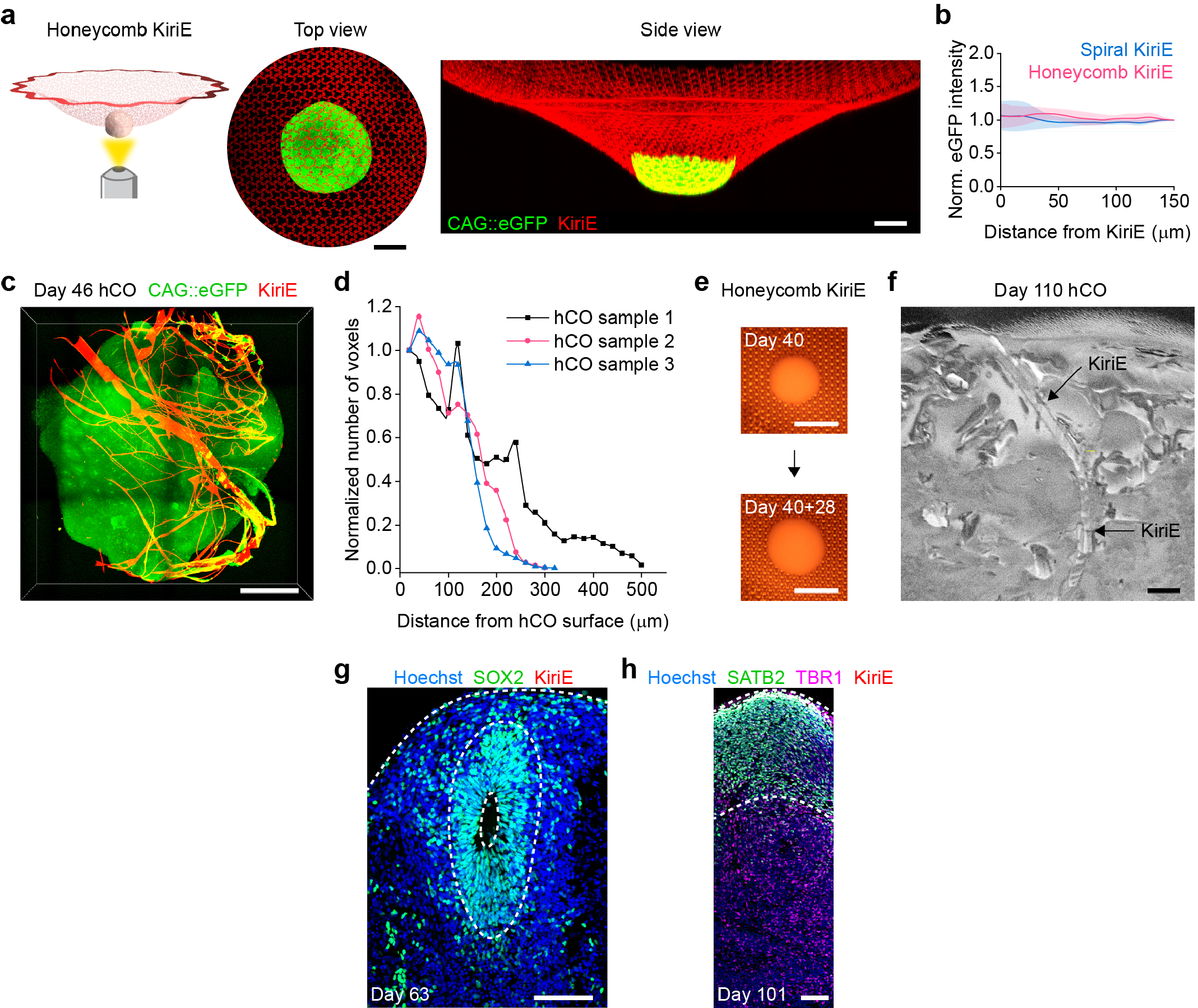
Integration of hCOs with KiriE. **a**, The top view and the side view of CAG::eGFP hCOs (green) on the honeycomb KiriE (red) reconstructed from live-cell confocal imaging. The level of the medium was lowered to better visualize the deformation of KiriE. Scale bars, 500 μm. **b**, Normalized fluorescence intensity (mean ± s.d.) of eGFP in the hCOs as a function of distance from spiral KiriEs (*n* = 3 hCOs) and honeycomb KiriEs (*n* = 3 hCOs), respectively. **c**, 3D interface of CAG::eGFP hCOs (green) and spiral KiriE (red) reconstructed from confocal imaging. The hCO-KiriE assembly was fixed and excised from the culture chamber before imaging. Scale bar, 500 μm. **d**, Normalized number of voxels of KiriE embedded within hCOs as a function of distance from the surface of hCOs after 26 days of integration. The KiriE is embedded as deep as 300–500 μm inside hCOs. **e**, Images of an hCO integrated with a honeycomb KiriE at different stages of differentiation. Scale bars, 1 mm. **f**, A FIB-SEM image showing the deformation of a KiriE line as it penetrates into the hCO. Scale bar, 2 μm. **g**, Immunostaining of SOX2^+^ neural progenitors in or near neural ventricular-like zones in KiriE-hCO cryosection. Scale bar, 100 μm. **h**, Immunostaining of deep layer (TBR1) and superficial layer (SATB2) cortical markers in KiriE-hCO cryosection. Scale bar, 100 μm.

**Extended Data Fig. 7.**
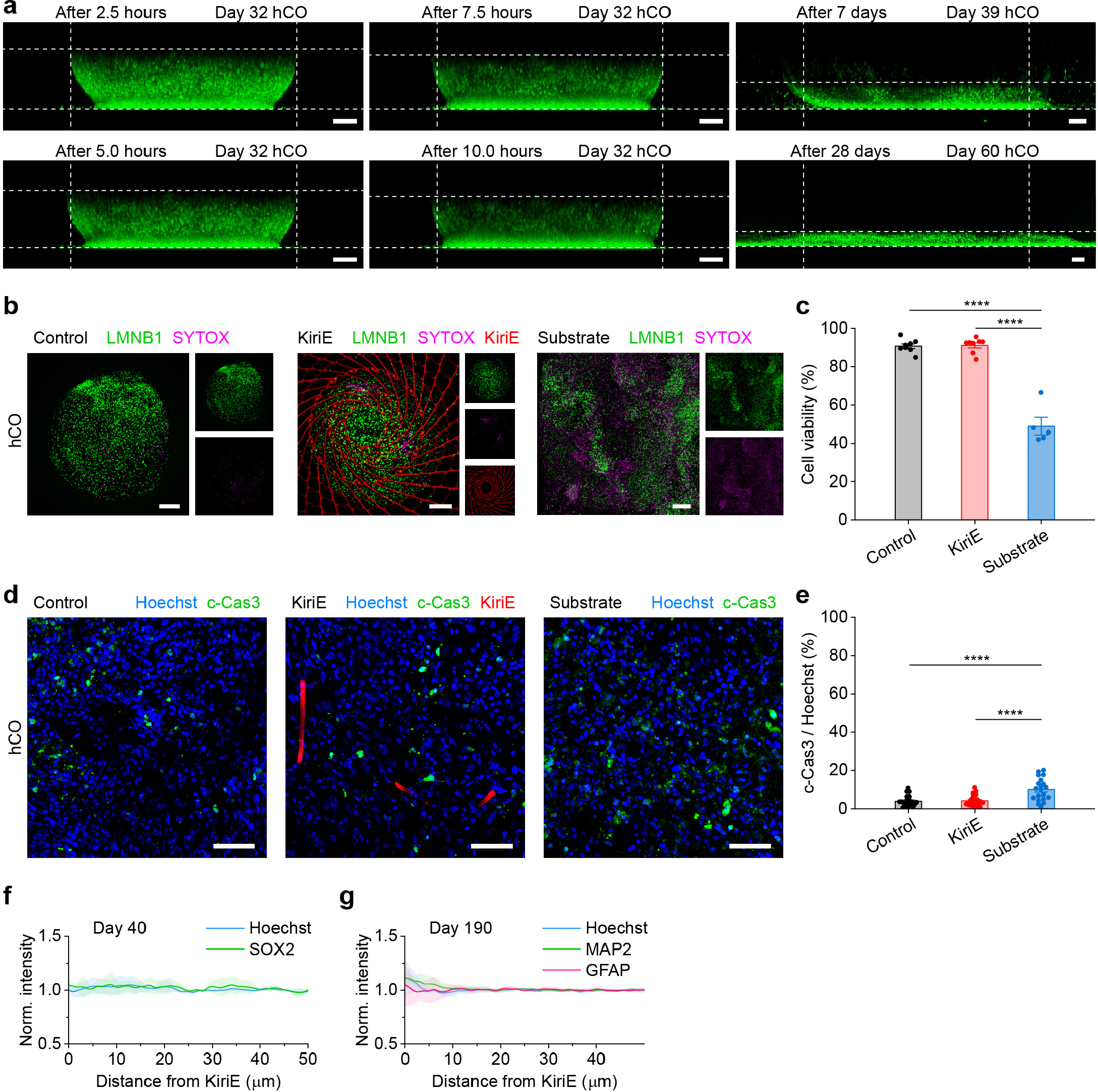
Morphological and cell survival characterizations of hCOs cultured in suspension (control), on KiriE, and on the adhesive substrate (Matrigel-coated glass-bottom plates). **a**, Time-lapse live-cell confocal images showing the flattening of a LMNB1-mEGFP hCO after being plated on an adhesive substrate. Scale bars, 100 μm. **b**, Live-cell confocal images using SYTOX™ deep red dead cell stain in day 95–109 LMNB1-mEGFP hCOs maintained in suspension (control), on KiriE, or on an adhesive substrate for 40–45 days. Scale bars, 200 μm. **c**, Quantification of cell viability across these conditions (*n* = 8 for control, *n* = 8 for KiriE, and *n* = 5 for adhesive substrate). One-way ANOVA test, *F*_2,18_ = 101.29, *P* < 1 × 10^−4^. *P* = 0.881 (control versus KiriE), \*\*\*\**P* < 1 × 10^−4^ (control versus adhesive substrate), \*\*\*\**P* < 1 × 10^−4^ (KiriE versus adhesive substrate). **d**, Immunostaining of c-Cas3^+^ cells in cryosections of day 91–102 hCOs maintained in suspension (control), on KiriE, and on the adhesive substrate for 41–48 days. Scale bars, 50 μm. **e**, Quantification of the ratio of c-Cas3 / Hoechst cells across these conditions (*n* = 20 for each group). One-way ANOVA test, *F*_2,57_ = 13.34, *P* < 1 × 10^−4^. *P* = 0.885 (control versus KiriE), \*\*\*\**P* < 1 × 10^−4^ (control versus adhesive substrate), \*\*\*\**P* < 1 × 10^−4^ (KiriE versus adhesive substrate). **f,g**, Normalized fluorescence intensity (mean ± s.d.) as a function of distance from KiriE for Hoechst and SOX2 in day 40 hCOs (*n* = 3 for **f**) and Hoechst, MAP2 and GFAP in day 190 hCOs (*n* = 4 for **g**) in KiriE-hCO cryosections shown in **Fig. 2i,j**. Data are presented as mean ± s.e.m. (**c,e**).

**Extended Data Fig. 8.**
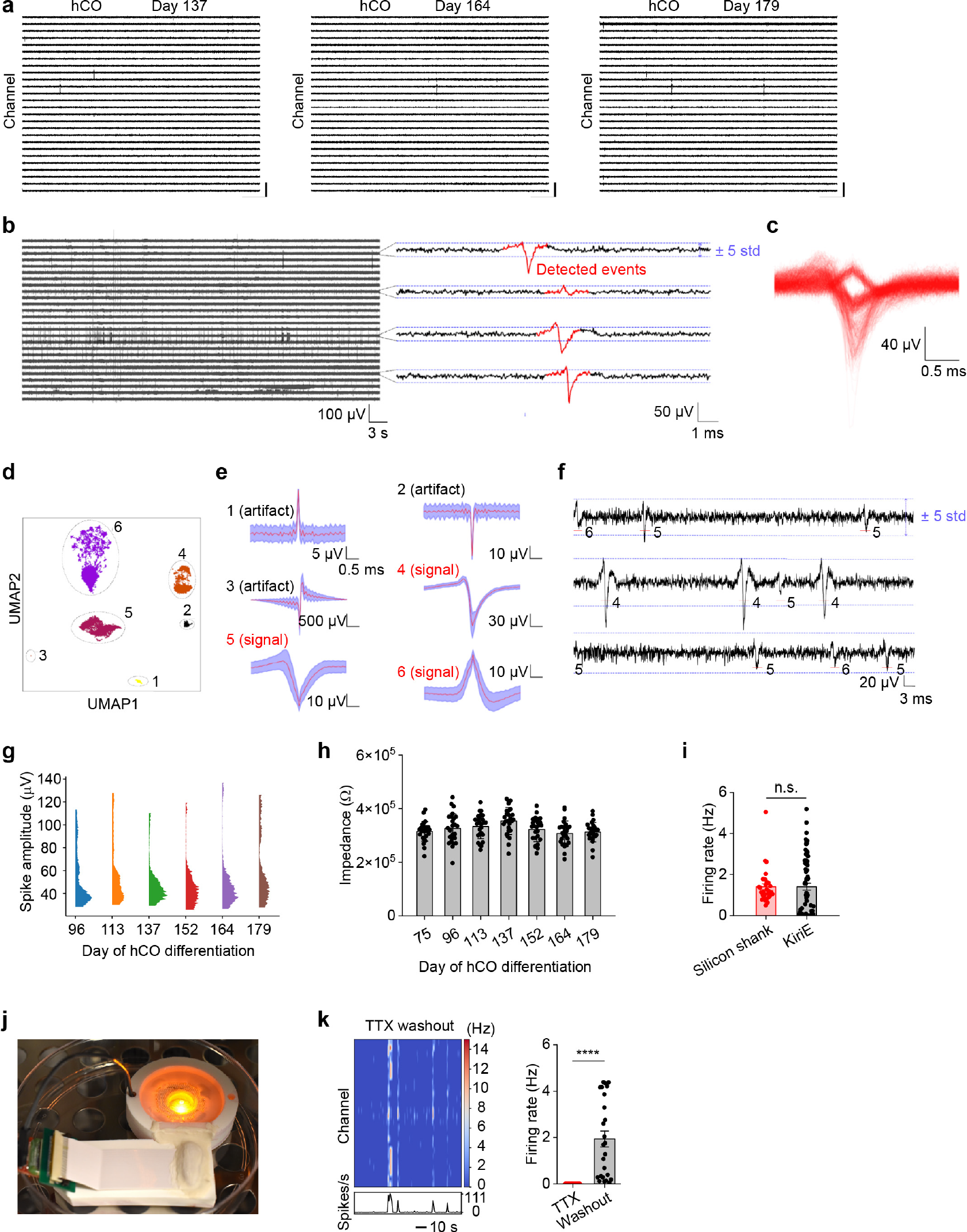
Electrophysiological recordings of hCOs and data analysis pipeline. **a**, Electrophysiological recordings of spontaneous activity of the hCO shown in **Fig. 3a** on days 137, 164, and 179. Scale bars, 100 ms (lateral) and 200 μV (vertical). **b**, Electrophysiological recordings sampled at 20 kHz are high-pass filtered at 150 Hz. For each channel, a sliding standard deviation with a 1 s window is computed. Any signal with a magnitude larger than 5 s.d. (dashed blue lines) is extracted (1.5 ms before and after the peak value for a total of 3 ms; marked in red) and stored. **c**, An overlay of all the detected signals shows a diversity of signal waveforms. **d**, UMAP dimensionality reduction is performed on every signal into two components. Density based clustering (DBSCAN) is performed on the component space. Each detected cluster corresponds to a given signal waveform. **e**, Signals from the numbered clusters in **d** are shown. The red line corresponds to the point-wise average of all signals within the detected clusters, and the blue shading corresponds to one point-wise s.d. Clusters 1 to 3 are clearly artifacts while clusters 4 to 6 are electrophysiological signals. The waveforms corresponding to neuronal signals (here clusters 4 to 6) are used as templates to find spikes of similar shape. **f**, Sample traces to illustrate how template matching enables the detection of small signals well within the initial 5 s.d. threshold shown in **b**. The red line indicates the position where a template was detected and the number corresponds to clusters in **e** and **e**. The template matching step is crucial in order to correctly estimate firing rates. **g**, Distributions of the spike amplitudes at different days of hCO differentiation. The average spike amplitudes across these days are 52.69, 56.82, 46.54, 45.49, 46.73, and 59.05 μV, respectively. **h**, Time-dependent electrode impedance at 1 kHz averaged over 26 electrodes in a KiriE-hCO assembly. Data are presented as mean ± s.d. **i**, Quantification of firing rates of day 116–125 hCOs measured by acutely inserted silicon shank electrodes and by KiriEs. Data are presented as mean ± s.e.m. **j**, A photograph of the optogenetic stimulation setup. **k**, The heatmap and the quantification of firing rates showing the recovery of neural activity upon TTX washout (*n* = 25 channels). \*\*\*\**P* < 1 × 10^-4^; two-tailed *t*-test. Data are presented as mean ± s.e.m.

**Extended Data Fig. 9.**
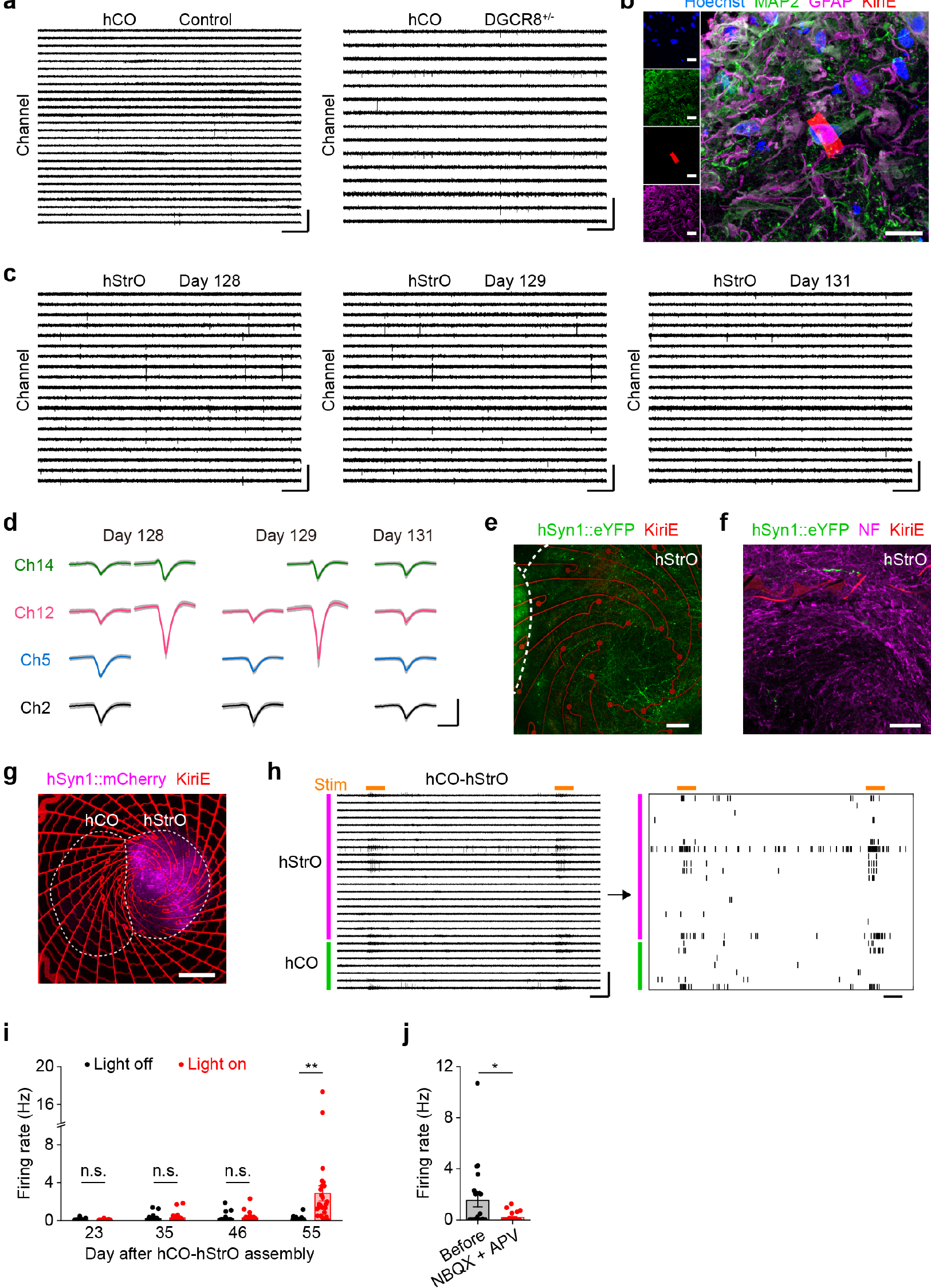
Electrophysiological recordings of hCOs, hStrOs, and cortico-striatal assembloids with integrated KiriE. **a**, Electrophysiological recordings of spontaneous activity of the DGCR8^+/-^ hCO and isogenic control hCO. Scale bars, 100 ms (lateral) and 200 μV (vertical). **b**, Immunostaining of MAP2^+^ neurons and GFAP^+^ glial lineage cells in KiriE-hStrO cryosections at day 271 of differentiation. Blue, green, magenta, and red colors represent Hoechst, MAP2, GFAP, and KiriE, respectively. Scale bar, 20 μm. **c**, Representative recordings of an hStrO on days 128, 129, and 131. Scale bars, 100 ms (lateral) and 200 μV (vertical). **d**, Single units from 4 representative channels in the hStrO on days 128, 129, and 131. Scale bars, 1 ms (lateral) and 50 μV (vertical). **e**, Live-cell confocal fluorescence image of a day 80+31 cortico-striatal assembloid on a KiriE. Scale bar, 100 μm. **f**, Immunostaining of neurofilament (NF) marker in cryosections of hCO-hStrO assembloids. Scale bar, 50 μm. **g**, Live-cell confocal fluorescence image of the mCherry channel of the cortico-striatal assembloid shown in **Fig. 4e**, showing that there are minimal projections from hStrO to hCO. Scale bar, 400 μm. **h**, Electrophysiological recordings and the corresponding raster plot of spiking activity during optogenetic stimulation of the cortico-striatal assembloid shown in **Fig. 4f**. The orange horizontal lines indicate light stimulation. Scale bars, 1 s (lateral) and 200 μV (vertical). **i**, Quantification of hStrO firing rates in the hCO-hStrO assembloid during the light-off (baseline) and the light-on (stimulation) phases at days 23, 35, 46, and 55 post-fusion (*n* = 27 channels over 18–30 light pulses). \*\**P* = 1.04 × 10^−3^; two-tailed *t*-test. Data are presented as mean ± s.e.m. **j**, Quantification of hStrO firing rates during the light-on phases in a cortico-striatal assembloid before and after the application of NBQX and APV (*n* = 22 channels analyzed over 18 light pulses). \**P* = 1.62 × 10^−2^; two-tailed *t*-test. Data are presented as mean ± s.e.m.

**Supplementary Table 1:**
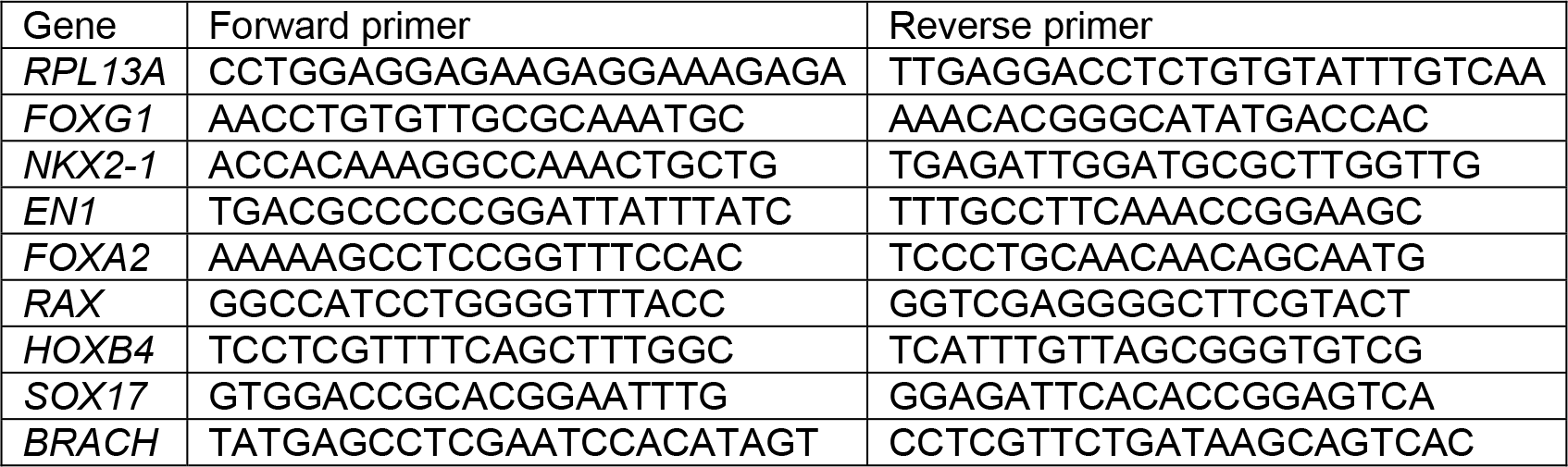
List of genes and primer sequences used for RT-qPCR.

**Supplementary Video 1:** An animation that shows the transformation of spiral KiriE from flat 2D pattern to 3D basket-like geometry and the integration with a neural organoid for long-term electrophysiology.

**Supplementary Video 2:** A Z-stack confocal image video of a 3D KiriE-hCO after 26 days of integration shows that KiriE is embedded within the hCO.

